# Efficient delivery of mRNA-LNPs in primary and secondary liver cancer

**DOI:** 10.1101/2025.03.18.643845

**Authors:** Laura J. Leighton, Yee Jing Gee, Sachithrani U. Madugalle, Maria Victorova, Nissa L. Carrodus, Kim R. Bridle, Sidney A. Howell, Xiaowen Liang, Gregory C. Miller, Chris L. D. McMillan, Danushka K. Wijesundara, David A. Muller, Darrell H. G. Crawford, Timothy R. Mercer, Seth W. Cheetham

**Affiliations:** Australian Institute for Bioengineering and Nanotechnology, The University of Queensland, Brisbane, QLD, Australia; BASE Facility, The University of Queensland, Brisbane, QLD, Australia; Faculty of Health, Medicine and Behavioural Sciences, The University of Queensland, Brisbane, QLD, Australia; Gallipoli Medical Research, Brisbane, QLD, Australia; The University of Queensland Frazer Institute, Brisbane, QLD, Australia; Envoi Pathology, Brisbane, QLD, Australia; School of Chemistry and Molecular Biosciences, The University of Queensland, Brisbane, QLD, Australia

## Abstract

Primary liver cancer is the sixth most prevalent cancer globally and is often diagnosed late, when treatment options are limited. Secondary liver cancer, arising from metastasis of other cancers to the liver, is a common complication of advanced solid cancers and a significant cause of cancer-related morbidity and mortality. Existing treatment options for advanced primary and secondary liver tumours have limited efficacy and new treatment modalities have the potential to improve patient outcomes. mRNA therapeutics are readily delivered to the healthy liver after systemic administration, but their uptake and expression within liver tumours is unclear. Here we show that intravenous delivery of mRNA-LNPs efficiently transfects virtually all hepatocytes in healthy, fibrotic, and cirrhotic liver, and also many cells of spontaneous hepatocellular carcinomas *in situ*. Delivery of mRNA is also possible to xenograft models of both primary and secondary liver cancer, albeit with attenuated protein expression relative to the normal liver. These findings demonstrate the potential for systemically delivered mRNA-LNP therapies for liver disease and cancer.

## Introduction

Primary liver cancer is the sixth most prevalent cancer worldwide, and the third-leading cause of cancer mortality^1^. Around 75% of primary liver cancers are hepatocellular carcinomas (HCC), which arise from malignant transformation of hepatocytes, the most common cell type in the liver. Primary liver cancer arises predominantly from chronic liver disease. Liver injury arising from chronic viral hepatitis, excessive alcohol use, and metabolic dysfunction-associated liver disease (MASLD) results in progressive fibrosis and ultimately cirrhosis, which predisposes individuals to HCC^2,3^. The increasing prevalence of MASLD in developed countries is driving increased incidence of primary liver cancer^3^.

When detected early, primary liver cancer can be cured by surgical resection or liver transplantation, but for patients with advanced disease, treatment options are limited and often burdensome (reviewed by Liu *et al*^4^). Transarterial chemoembolization (TACE) and selective internal radiation therapy (SIRT) deliver therapeutics directly to the tumour while also interrupting its blood supply^5–9^. Systemic drug therapies are typically used in advanced HCC, in particular sorafenib and lenvatinib (inhibitors of protein kinases including VEGFR), and the combination of atezolizumab (a PD-L1 inhibitor) and bevacizumab (which targets VEGF-A)^9–12^. These systemic treatments can extend survival for several months on average, but are rarely curative, and associated with burdensome side effects including cutaneous reactions, stomatitis, peripheral neuropathy, gastrointestinal symptoms, and fatigue. The prognosis for primary liver cancer remains poor, with 5-year survival of approximately 22%^13,14^.

Secondary liver cancer, caused by metastasis of other cancers to the liver, is a substantial contributor to cancer morbidity and mortality. Around 50% of patients with metastatic colorectal, breast and pancreatic cancer, and around 35% of patients with metastatic lung cancer, develop liver metastases^15–18^, and these liver tumours are a frequent cause of cancer death^15,17–20^. For secondary liver cancer patients also, treatment options are limited and include liver resection, ablative techniques, embolization therapies, and systemic chemotherapy or immunotherapy^21–23^. However, secondary liver cancer generally carries a poor prognosis^20,23^. Alternative treatment modalities are urgently needed for primary and secondary liver cancers.

The rapid development of SARS-CoV-2 vaccines demonstrated powerfully the safety and efficacy of synthetic messenger RNA (mRNA) medicines. Synthetic mRNAs are comprised of a coding sequence which can encode any therapeutic protein, flanked by 5′ and 3′ untranslated regions from highly-expressed and stable mammalian mRNAs, and a poly(A) tail. To attenuate the innate response to foreign RNA^24^, mRNA drugs are depleted of secondary structures and uridine residues, and remaining uridines substituted with the uridine analogue N1-methylpseudouridine. Formulation of mRNA into lipid nanoparticles protects the mRNA and enables delivery to cells via endocytosis. A small percentage of the delivered mRNA escapes the endosome into the cytoplasm, where it is then translated by cytoplasmic ribosomes to produce the target protein. As of 2023, over 13 billion doses of mRNA vaccines had been administered globally with an adverse-event rate comparable to other vaccines^25^, demonstrating the fundamental effectiveness and safety of this drug modality.

Intravenously injected mRNA-LNPs are efficiently internalised in the liver, where the mRNA-encoded protein is expressed by hepatocytes for several days^26^. Hepatic mRNA uptake encompasses virtually all hepatocytes, and subpopulations of endothelial and Kupffer cells^27^. The natural tissue tropism of LNPs for hepatocytes has enabled experimental mRNA-based treatments of genetic metabolic disorders including methylmalonic acidemia^28^, acute intermittent porphyria^29^ and Fabry disease^30^ in animal models, and a systemically administered mRNA-LNP treatment for propionic acidemia has shown promise in early human trials^31^. Importantly, these studies have demonstrated the safety and efficacy of repeated mRNA-LNP administration, which is critical if mRNA drugs are to be used for ongoing treatment of liver disease.

The efficient delivery of mRNA-LNPs to healthy livers raises the possibility of using mRNA therapeutics for the treatment of liver disease and cancer. Primary liver cancer typically arises on a background of chronic liver disease and hepatic fibrogenesis. One study reported good expression of mRNA-LNPs in hepatocytes in several mouse models of fibrosis and cirrhosis, and a therapeutic effect of *Hnf4a* mRNA against fibrosis^32^. However, the extent of mRNA-LNP delivery into liver tumours has never been rigorously assessed.

Here we show that following intravenous administration, mRNA-LNPs enable efficient protein expression not only in healthy liver, but also in fibrotic and cirrhotic liver, spontaneous hepatocellular carcinomas *in situ*, and xenograft models of primary and secondary liver cancer in mice. These findings suggest that intravenous injection of mRNA-LNPs without a specific targeting moiety is a promising means to deliver therapeutic proteins for the treatment of liver disease and tumours.

## Results

### Efficient delivery of mRNA-LNPs to the healthy mouse liver

We initially set out to establish a baseline for mRNA-LNP delivery to and expression in the mouse liver. We selected a formulation for mRNA-LNPs based on the ionisable lipid SM-102, which is approved for clinical use. To assess the delivery of mRNA to the liver, we intravenously injected adult male mice (n=5) with mRNA-LNPs encoding enhanced green fluorescent protein (eGFP) (**Figure 1a**). Strong and specific eGFP fluorescence was observed in the liver after 24 hours via *ex vivo* fluorescence imaging, with measured fluorescence values around 10-fold higher than background (tissue autofluorescence) (**Figure 1b,c**). Background-subtracted fluorescence (radiant efficiency) of whole livers was significantly different between mRNA-injected and uninjected mice (Student’s t-test, n=4-5, p<0.0001) (**Figure 1d**). Western blotting detected eGFP protein strongly from the liver tissue of these mice (**Figure 1e**). Virtually all hepatocytes took up eGFP mRNA as evident from immunohistochemistry (**Figure 1f, Supplementary Figure 1**), with variable staining intensity between cells, and no obvious bias in spatial distribution. This demonstrated the efficient uptake and expression of mRNA by the healthy liver, and provided a basis for comparison of the results in diseased liver.

**Figure 1:**
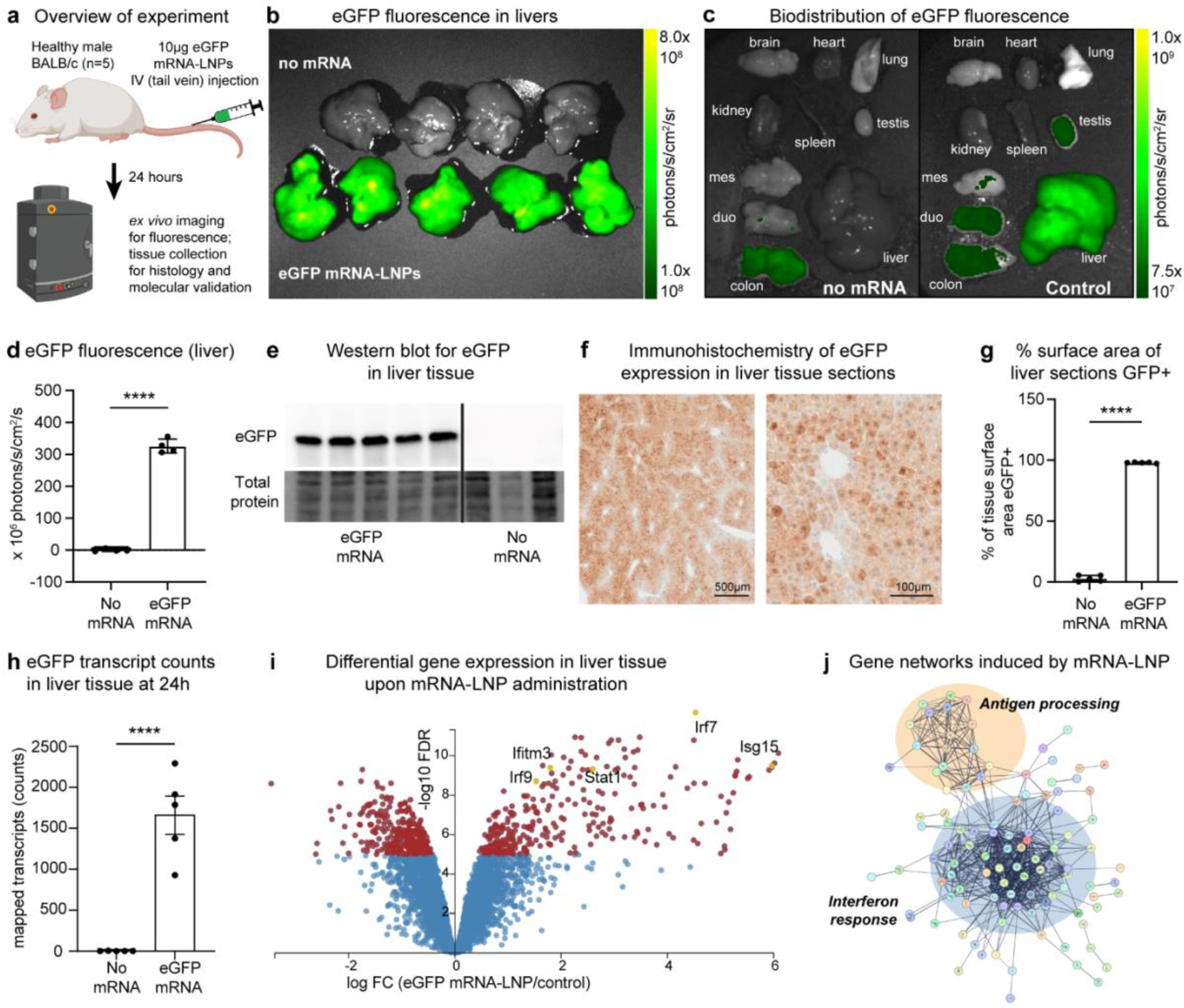
Efficient delivery of mRNA-LNPs to the healthy mouse liver. (**a**)We injected 5 healthy mice with 10µg eGFP mRNA-LNPs via the tail vein and investigated reporter expression after 24 hours. **(b)** Strong eGFP fluorescence was visualised in all injected livers (bottom row). **(c)** *ex vivo* fluorescence imaging of organs from injected and uninjected mice shows specificity of eGFP expression in the liver. Fluorescence scales were set to eliminate autofluorescent signal in the uninjected liver; autofluorescent signal is visualised in the colon and duodenum of both the injected and uninjected mouse. **(d)** Quantitation of background-subtracted eGFP fluorescence confirms the strong and specific detection of eGFP in the livers of eGFP-injected mice (Student’s t-test, n=4-5, t=29.33(6), p<0.0001). Error bars represent mean ± SEM. **(e)** Western blotting detects a strong band corresponding to eGFP in the livers of mice injected with eGFP mRNA-LNPs; no band is detected in liver tissue of uninjected mice. **(f)** Immunohistochemistry for eGFP in liver tissue sections from injected mice shows strong eGFP staining in virtually all hepatocytes. There is moderate variation in staining intensity between hepatocytes. **(g)** Quantification of GFP-positive surface area for IHC sections in GFP-injected mice shows extremely high coverage of liver tissue, with very low background in uninjected mice (Welch’s t-test, n=5, t=84.81(4.204), p<0.0001). Error bars represent mean ± SEM. **(h)** eGFP transcripts were readily detected in mouse liver tissue 24 hours after injection, with mapped transcript counts between approximately 1000 and 2300; this was significantly different from control with an average of one count (Mann-Whitney test, n=5, U=0, p=0.0079). **(i)** 624 genes were differentially expressed in the liver between mRNA-LNP injected and saline-injected mice, with the significance threshold of 5 * –log10 FDR. These included interferon response factors (IRF7, IRF9, IFITM3, ISG15) and the cytokine-responsive transcription factor STAT1. **(j)** Gene networks induced in the liver by mRNA-LNP injection; upregulated genes cluster into antigen-processing and interferon-response networks.

To quantify the distribution of eGFP expression in tissue sections, we trained a machine learning algorithm, *StainDetectAI*, to distinguish stained from unstained liver tissue in IHC microscopy images (**Supplementary Figure 2**). The model was trained using a supervised machine learning approach, using a training dataset consisting of manually annotated IHC images of GFP-positive and GFP-negative liver tissue. The model has specificity of 98% and sensitivity of 92%, with an overall prediction accuracy of 95%. Using *StainDetectAI*, we showed that eGFP mRNA-LNP injected animals had around 97% of liver section surface area stained for eGFP (**Figure 1g**), while on average less than 3% of surface area was falsely detected as stained in liver sections from uninjected mice. The surface area detected as stained between injected and uninjected mice was significantly different (Welch’s t-test, n=5, p<0.0001).

An important consideration for the safety and efficacy of mRNA-LNP therapeutics delivered via the intravenous route is the induction of an immune response, and potentially other off-target gene expression, in the liver. To determine whether the mRNA-LNP platform impacts on liver gene expression, we sequenced RNA from liver tissue of healthy mice injected with eGFP mRNA-LNPs (n=5) and saline-injected controls (n=5) 24 hours after injection. We detected abundant eGFP transcripts (averaging 1661 counts) in the livers of mRNA-injected mice, with negligible background (average 1 count in uninjected mice), demonstrating that mRNA can persist in liver tissue for at least a day after injection (**Figure 1h**); the difference was significant (Mann-Whitney test, n=5, p=0.0079). Consistent with the known immunogenicity of mRNA-LNPs, intense upregulation of genes involved in innate immunity was observed in the liver tissue of mRNA-injected mice relative to uninjected controls, including IRF7/9, STAT1 and IFITM3. In total, 295 genes were significantly upregulated and 323 genes significantly downregulated (**Figure 1i, Supplementary Table 1, Supplementary Figure 3**). Notably, PD-L1 (CD274) was significantly upregulated in liver cells following mRNA-LNP treatment (Welch’s t-test, n=5, p<0.0001), substantiating recent reports that mRNA-LNP administration can increase checkpoint ligand expression. The widespread immune response suggests that the mRNA-LNP platform is inherently inflammatory in the liver and that the evaluation of candidate therapeutics must be carefully controlled to account for platform effects.

### Delivery of mRNA-LNPs in liver fibrosis and cirrhosis

We next investigated delivery of mRNA-LNPs in a physiologically-relevant mouse model of progressive liver injury. *Mdr2* knockout mice are unable to secrete phospholipids into the bile, resulting in the development of sclerosing cholangitis with severe liver fibrosis by early adulthood and spontaneous hepatocellular carcinoma by 1 year of age^37,38^. We used the *Mdr2* knockout model to evaluate mRNA-LNP delivery to the liver in the context of liver fibrosis and cirrhosis (**Figure 2a**). We injected eGFP mRNA-LNPs intravenously into 4 male *Mdr2* knockout mice at 11 months of age, and 14 hours post-injection, we observed fluorescent signal 3-4 fold above background in the liver tissue, albeit attenuated 2-fold relative to healthy control (**Figure 2b**). Biodistribution of eGFP fluorescence in *Mdr2* knockout mice was comparable to that observed in healthy BALB/c mice (**Figure 2c**), indicating that off-target expression of the mRNA in other organs and tissues is not increased in the presence of liver injury.

**Figure 2:**
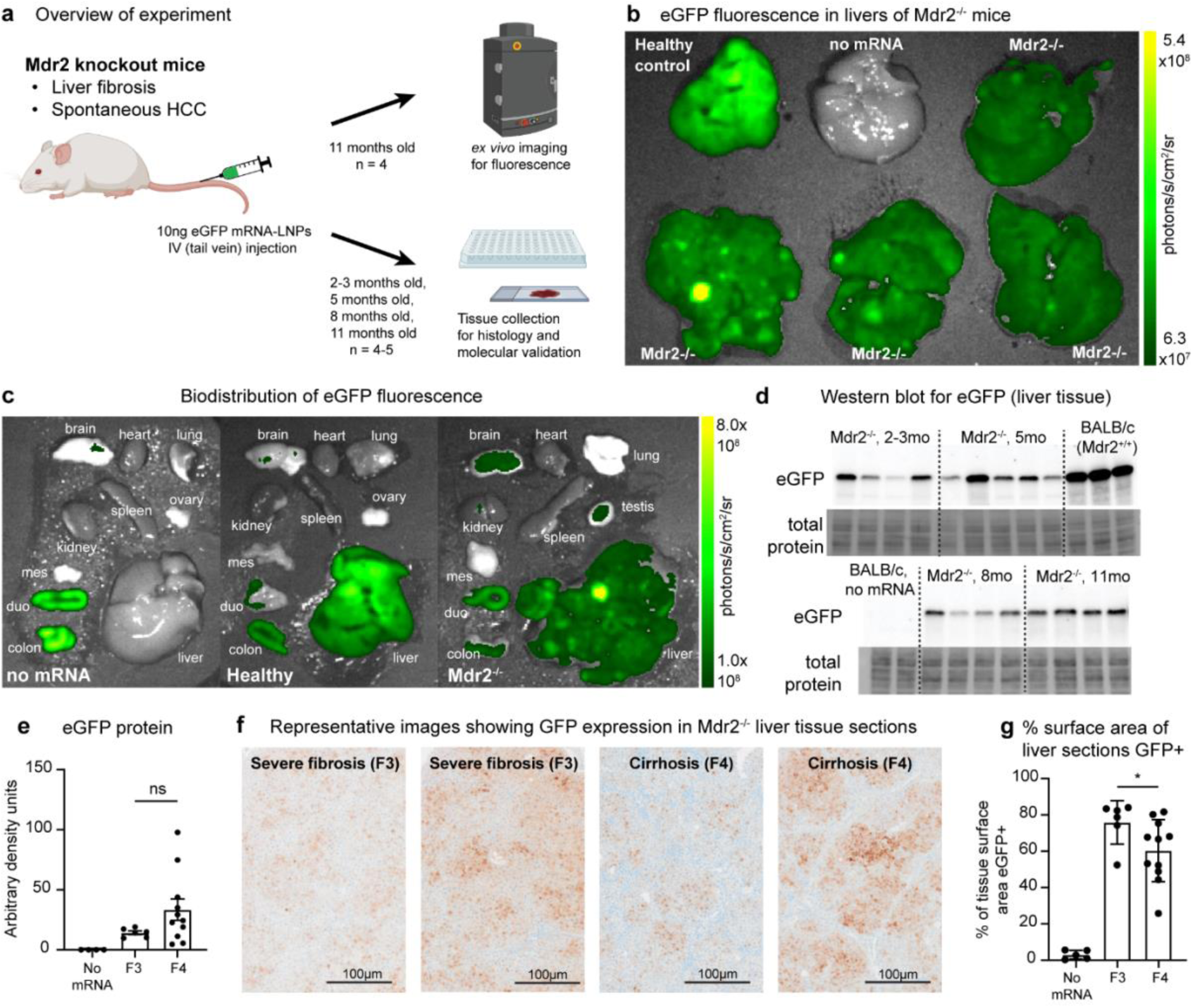
Delivery of mRNA-LNPs in liver fibrosis and cirrhosis. (**a**) We injected *Mdr2*^−/−^ mice with 10µg eGFP mRNA-LNPs via the tail vein and investigated reporter expression. **(b)** Moderately strong eGFP fluorescence was evident in all injected livers; the fluorescence intensity was reduced compared to a healthy control injected with the same mRNA-LNPs at the same time. **(c)** *ex vivo* fluorescence imaging of organs from a GFP-injected *Mdr2*^−/−^ mouse shows that attenuation of mRNA-LNP delivery to the liver does not result in broadly altered biodistribution of eGFP protein expression. Autofluorescent signal was detected from the brain and gut tissues of all mice including healthy and uninjected controls. **(d)** Western blotting detects eGFP from the liver tissue of 17 *Mdr2*^−/−^ mice, with substantial variation between individuals in the quantity of eGFP protein, and reduced protein abundance relative to healthy BALB/c mice. **(e)** Quantification of eGFP protein expression from Western blotting showing greater variability in protein expression for *Mdr2*^−/−^ mice with a fibrosis score of F4 (cirrhosis) relative to F3 (severe fibrosis). The total protein abundance in F4 mice trended higher but this difference was not statistically significant (Welch’s t-test, n=6-11, t=2.162(10.58), p=0.0545). Error bars represent mean ± SEM. **(f)** Representative immunohistochemistry results from *Mdr2*^−/−^ mice show the range of possible outcomes for eGFP expression. **(g)** Analysis of IHC images with *StainDetectAI* found that mice with severe fibrosis (F3) had detectable eGFP staining across a higher surface area compared to mice with cirrhosis (F4) (Mann-Whitney test, n=6-11, U=12, p=0.0365). Error bars represent mean ± SEM.

Separately, eGFP mRNA-LNPs were injected into male *Mdr2*^−/−^ mice aged 2-3 months (n=4), 5 months (n=4), 8 months (n=5) and 11 months (n=4). Using Western blotting, we found that eGFP protein was evident in liver tissue from all mice after 24h, with substantial variation between individuals; protein density ranged from 4.65 to 97.85 (arbitrary units, **Figure 2d,e**). Transfection of hepatocytes was widespread in most livers (**Figure 2f, Supplementary Figure 4**). However, the staining intensity and the proportion of transfected cells were both reduced relative to healthy livers (shown in **Figure 1f**), and staining was marginal to absent within fibrotic bands, demonstrating that extensive fibrosis reduces but does not ablate mRNA-LNP delivery efficiency. Using *StainDetectAI,* we found that the total surface area of liver tissue positive for GFP staining (quantified using *StainDetectAI)* ranged from 26% to 84% across all 17 mice.

Given the variability in disease severity between individual mice, we next evaluated eGFP mRNA expression relative to the severity of liver disease. We used the METAVIR scoring system to assess the extent of liver fibrosis based on H&E stained liver tissue sections (**Supplementary Figure 4**); out of 17 mice, 6 received a score of F3 (severe fibrosis) and 11 were scored F4 (cirrhosis). Importantly, we did not find a significant difference in eGFP protein expression between animals scoring F3 and F4 (Welch’s t-test, n=6-11, p=0.0545) (**Figure 2e**). When comparing the result from *StainDetectAI* between F3 and F4 animals, we found that the stained surface area was significantly negatively associated with the degree of liver fibrosis (**Figure 2g**); animals with a fibrosis score of F3 averaged 76% surface area staining, while F4 animals averaged 60% (Mann-Whitney test, n=6-11, p=0.0365). One possible explanation for this result is that mice with more severe disease have more of the liver section surface area occupied by fibrotic bands, resulting in reduced hepatocyte surface area. We therefore quantified the stained surface area in 10 randomly-selected liver tiles per animal from which any fibrotic areas were manually masked. This analysis found that in these hepatocyte-only samples, there remained a significant difference between animals with a fibrosis score of F3 which averaged 83% surface area staining, and animals scoring F4 which averaged 67% (Mann-Whitney test, n=6-11, p=0.0103) (**Supplementary Figure 5**). This shows that the difference in distribution of mRNA expression between F3 and F4 animals is not merely a function of the increased surface area of fibrotic bands.

Overall, these results show that while fibrosis and cirrhosis reduce the efficiency of mRNA-LNP delivery to the liver, robust and widespread mRNA delivery is still evident, with the majority of hepatocytes transfected.

### Delivery of mRNA-LNPs to spontaneous hepatocellular carcinomas *in situ*

*Mdr2* knockout mice develop spontaneous hepatocellular carcinomas with age, mirroring disease progression in humans. Tumours are usually observed by 6-8 months of age, with males older than 10 months typically developing multiple large HCCs. The *Mdr2* knockout mouse is a physiologically relevant model for human HCC because, similar to humans with chronic liver disease, HCCs in these mice arise from malignant transformation of injured hepatocytes on a background of progressive fibrosis or cirrhosis.

To investigate the efficacy of mRNA-LNP delivery in a physiologically relevant HCC model, we first compared eGFP expression between liver tissue and liver tumours of 11 month old male *Mdr2*^−/−^ mice using *ex vivo* fluorescence imaging. Notably, many liver tumours were evident on the external surfaces of the livers (**Supplementary Figure 6**), including several large tumours with increased fluorescence relative to the surrounding liver tissue (**Figure 2b**, **Figure 3a**). The average fluorescence of the tumours (262.6 x 10^6^ photons/s/cm^2^/sr) was not significantly different from that of the liver tissue (97.1 x 10^6^ photons/s/cm^2^/sr) (Mann-Whitney test, n=4-77, p=0.8738) (**Figure 3b, Supplementary Figure 7**). There was no correlation between tumour size and fluorescence intensity (simple linear regression, R^2^=0.0661) (**Figure 3c**).

**Figure 3:**
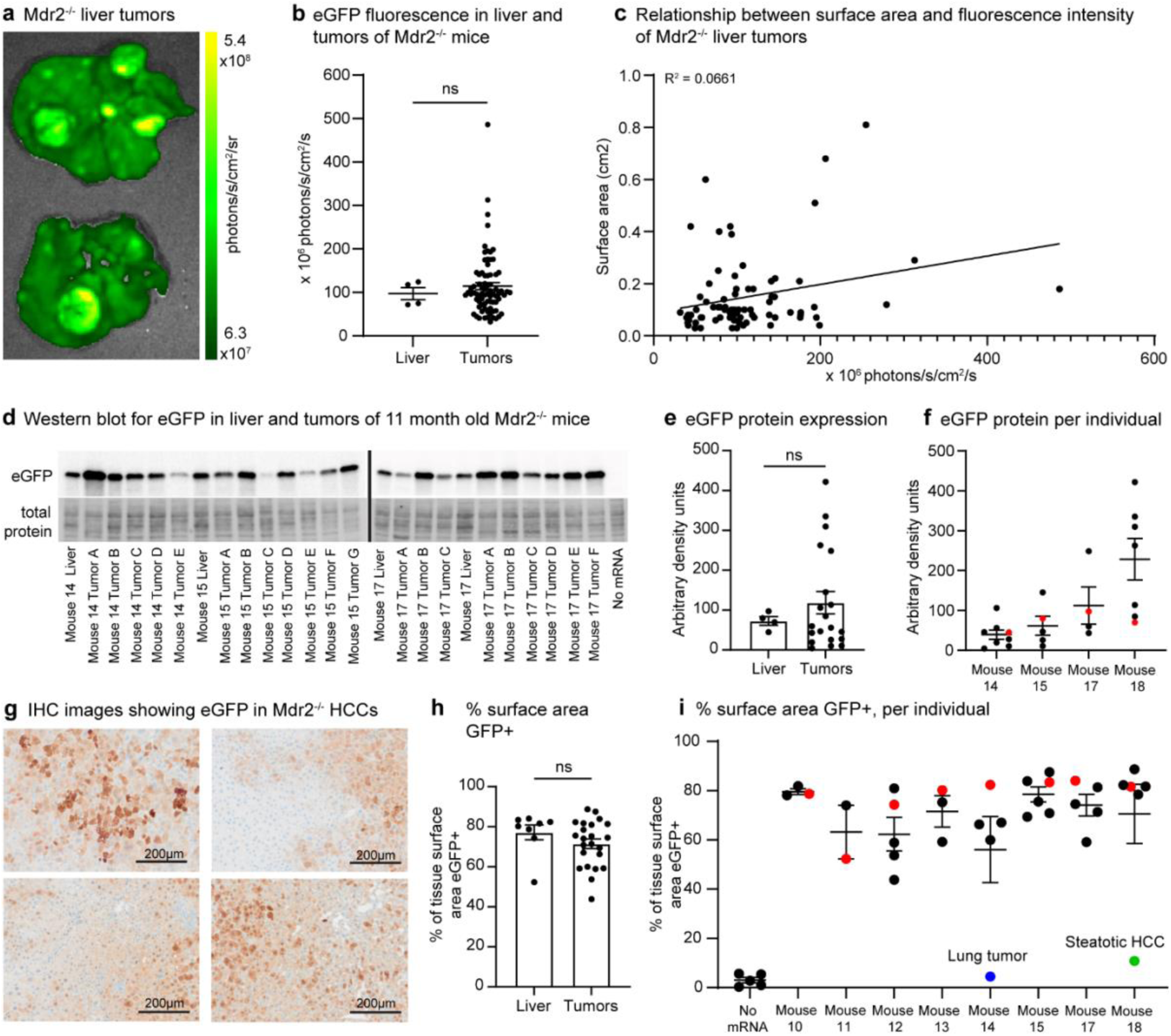
Delivery of mRNA-LNPs to spontaneous hepatocellular carcinomas *in situ*. (**a**) Fluorescence image showing large HCCs on the livers of *Mdr2*^−/−^ mice. **(b)** Plot of fluorescence intensity of whole livers (n=4) and individual tumours visible on the external surfaces of the same livers (n=77). There was no significant difference in fluorescence intensity between livers and tumours (Mann-Whitney test, n=4-77, U=146, p=0.8738). Error bars represent mean ± SEM. **(c)** There is no correlation between the external surface area of liver tumours and the fluorescence intensity (simple linear regression, F(1,75)=5.308, p=0.0240, R^2^=0.06609). **(d)** Western blotting detects eGFP from livers and tumours of 4 *Mdr2*^−/−^ mice 11 months of age. The quantity of eGFP protein detected from tumours is broadly comparable to that detected from livers, and also shows variation. **(e)** Quantitation of eGFP protein expression determined by Western blotting, showing greater variability in protein expression in tumours (n=21) than in liver tissue (n=4). The abundance of eGFP protein in tumours was not significantly different from liver tissue (Welch’s t-test, n=4-21, t=1.519(21.99), p=0.1430). Error bars represent mean ± SEM. **(f)** Quantitation of eGFP protein separated by animal; the red point in each column represents the eGFP abundance in liver tissue, and the black points the tumours from the same animal. Error bars represent mean ± SEM. **(g)** Representative IHC images show a range of possible outcomes for eGFP expression in tumour tissue, including broadly good delivery, variability in staining intensity between adjacent cells and between different regions of the image, and borders between stained and unstained regions of the tumour which are otherwise similar in appearance. **(h)** Analysis of IHC images with StainDetect AI shows that the surface area of tissue sections with detectable IHC staining is not significantly different between liver tissue and tumours (Mann-Whitney test, n=8-25, U=60, p=0.1580). Error bars represent mean ± SEM. **(i)** Quantification of IHC surface area positive for eGFP separated by animal; the red point in each column represents the stained surface area in liver tissue, and the black points the tumours from the same animal. Note poor delivery to a lung adenocarcinoma (blue; eGFP not detected above background level) and a steatotic HCC (green; eGFP detection marginal.)

Using Western blotting, we found that expression of eGFP protein in tumours was comparable to the expression in liver tissue, with considerable variability in protein abundance observed between individual tumours (arbitrary protein density units, tumours range 4.3-774.9 and average 150.1, liver tissue range 44.9-97.1 and average 72.9) (**Figure 3d-f**). There was no significant difference between the average amount of eGFP protein detected in tumours relative to the liver tissue (Welch’s t-test, n=4-21, p=0.1430) (**Figure 3e**).

To investigate the pattern of eGFP expression at a cellular level, immunohistochemistry was performed on sections of multiple individual tumours per mouse. Immunohistochemistry for eGFP in tumour sections confirms that eGFP expression is variable both within tumours (with all tumours including cells both lightly and intensely stained for eGFP) and between tumours (with some tumours showing more stained cells and/or more intense staining than others) (**Figure 3g, Supplementary Figure 8**). We again used our custom model *StainDetectAI* to determine the percentage of tumour section surface area that exhibited positive GFP staining, and found that surface area staining of spontaneous HCCs ranged from 43.7% to 88.7% (**Figure 3h,i**). There was no significant difference in the percentage of surface area stained between liver tissue and tumours (Mann-Whitney test, n=8-25, p=0.1580) (**Figure 3h**). Notably, tumour C from animal 18 was the only steatotic hepatocellular carcinoma in the dataset, and eGFP expression in this tumour was almost undetectable via both Western blotting and immunohistochemistry (**Figure 3d,i, Supplementary Figure 9**). Additionally, a spontaneous adenocarcinoma of the lung was incidentally recovered from one of the 11 month old animals. This tumour was also analysed with *StainDetectAI*, and eGFP staining was not detected above background levels (**Figure 3i, Supplementary Figure 10**). These findings demonstrate that intravenous delivery of mRNA-LNPs results in specific expression within most hepatocytes and many cells within spontaneously-occurring hepatocellular carcinomas *in situ*, with minimal delivery to other organs and tissues, likely including other cancers. This restricted delivery pattern suggests that mRNA-LNP therapeutics for HCC could be administered systemically with limited extra-hepatic off-target activity.

### Delivery of mRNA-LNPs to a xenograft model of primary liver cancer

Animals bearing tumour xenografts derived from human cancer cell lines are a commonly-used model system for testing new anti-cancer therapeutics. To determine whether mRNA-LNPs are delivered to liver tumour xenografts as in spontaneously-occurring HCCs, we established an orthotopic xenograft model using HuH-7 human hepatocellular carcinoma cells modified to constitutively express mCherry and firefly luciferase (**Supplementary Figure 11**). Cells were engrafted into the livers of thioacetamide-treated BALB/c nude mice by single direct injection. Due to the aggressive growth of the HuH-7 tumours, mice were injected with eGFP mRNA-LNPs every third day beginning after a 1-week tumour establishment period, and when estimated tumour size reached 1cc, mice were euthanised 24 hours after the most recent eGFP mRNA-LNP injection (**Figure 4a**).

**Figure 4:**
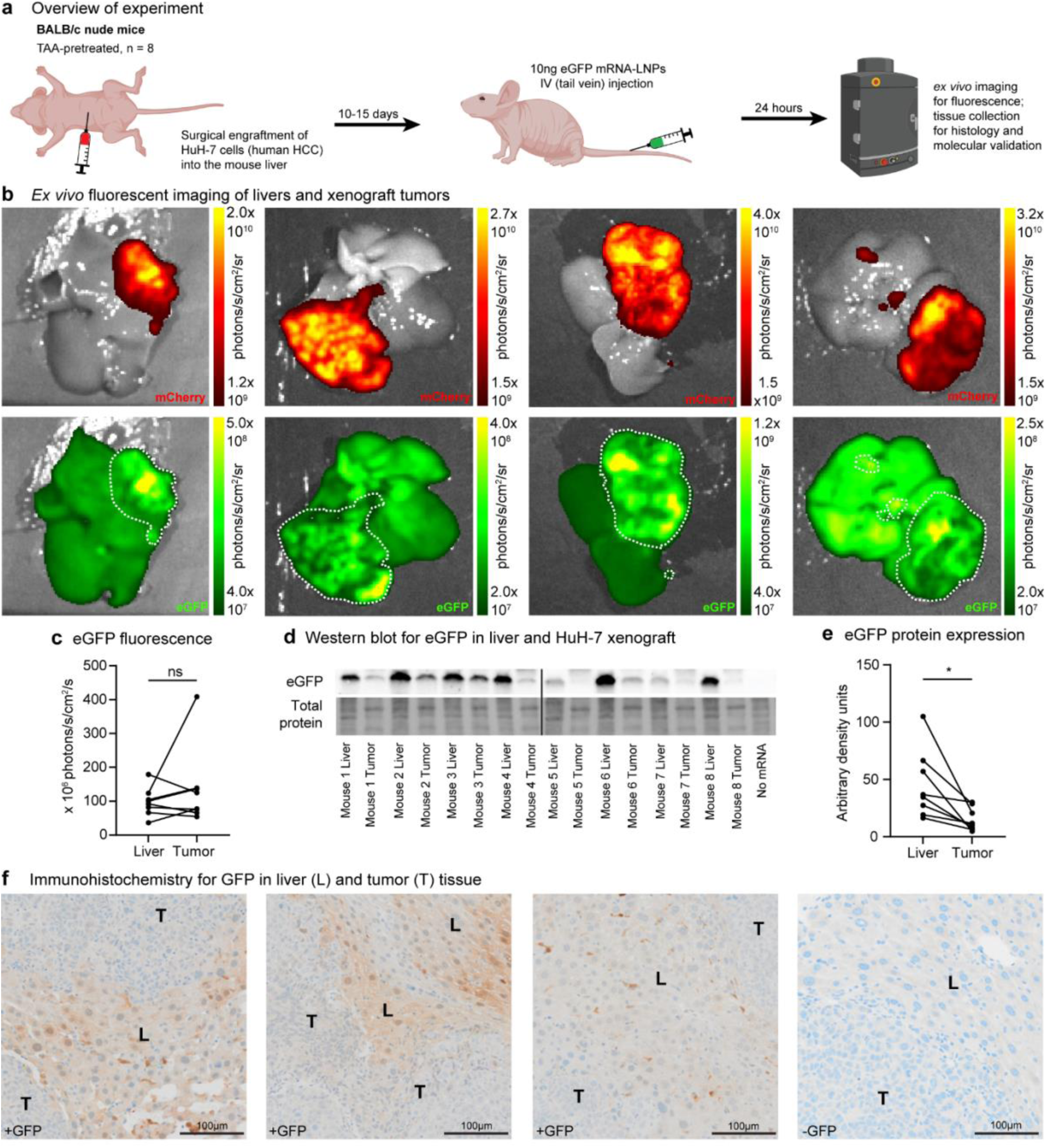
Delivery of mRNA-LNPs to a xenograft model of primary liver cancer. (**a**) We surgically engrafted HuH-7 human HCC cells which constitutively express mCherry into the livers of BALB/c nude mice. Tissues were analysed approximately 24 hours after an injection of 10µg eGFP mRNA-LNPs via the tail vein. **(b)** *Ex vivo* fluorescent imaging of entire livers with xenograft-derived tumours shows strong eGFP fluorescence in the liver tissue and also the tumours, marked by expression of mCherry. **(c)** Quantitation of eGFP fluorescence confirms broadly comparable signal between tumours and non-tumour liver tissue, with no significant difference found between liver and tumour (Paired t-test, n=8, t=1.002(7), p=0.3496). **(d)** Western blotting detects eGFP from all livers and tumours, with noticeably higher signal in the liver tissue, and some bands from tumour detected very faintly. **(e)** Quantitation of eGFP protein expression determined by Western blotting, showing lower signal in liver than tumour of each animal (Paired t-test, n=8, t=3.168(7), p=0.0158). **(f)** Representative immunohistochemistry images of liver and tumour tissue from 3 different animals show clear GFP staining in HuH-7 derived tumours (T), and staining of a higher intensity in adjacent liver tissue (L). An image of HuH-7 derived tumour and adjacent liver tissue from a mouse not injected with eGFP and stained with anti-GFP antibody is provided for comparison.

To investigate the delivery of mRNA-LNPs to the tumours, we first used *ex vivo* fluorescent imaging of the eGFP reporter. Fluorescent signal well above background level was observed in the livers (range 36.4-179.1, average 97.9 x 10^6^ photons/s/cm^2^/sr) and liver tumours (range 54.6-408.7, average 135.4 x 10^6^ photons/s/cm^2^/sr) of all mice (n=8) (**Figure 4b,c**). There was no significant difference in fluorescent signal between liver tissue and tumours (Paired t-test, n=8, p=0.3496). We considered the possibility that residual healthy liver tissue overlying the tumours might uptake mRNA-LNPs and express eGFP, resulting in high tumour-surface fluorescence values which may not represent the uptake and expression of mRNA within the tumour interior. Therefore, to investigate penetration of mRNA-LNPs into the tumour interior, we performed fluorescence imaging on cross-sections of several tumours and observed mostly uniform fluorescence intensity within the tumours (**Supplementary Figure 12**). We also expected that mRNA-LNP penetration might be better in small tumours than in large tumours, but we found that there was no correlation between tumour size (measured by area of the region of interest drawn around the tumour) and the fluorescence intensity (simple linear regression, R^2^=0.0002) (**Supplementary Figure 13**). Surprisingly, there was also no correlation between the fluorescence intensity measured from the liver and tumour from the same animal (simple linear regression, R^2^=0.1620) (**Supplementary Figure 13**).

Western blotting supported the findings from fluorescence imaging: a clear eGFP band was detected from tumour samples, but with variability in band intensity (**Figure 4d**), with bands from two tumours detected extremely faintly. Protein expression was significantly lower in the tumour samples than in the liver tissue (range 16.3-104.9 and average 45.3 in the liver and range 4.7-30.0 and average 14.9 in the tumours) (Paired t-test, n=8, p=0.0158) (**Figure 4e**). Immunohistochemistry for eGFP in 3 tumours showed weak expression of eGFP in virtually all tumour cells, with some cells at the margins of most tumours showing higher signal (**Figure 4f, Supplementary Figure 14**). Expression of eGFP was substantially weaker in the HuH-7 tumours relative to the adjacent liver tissue. These results demonstrate that it is possible to deliver mRNA-LNPs to orthotopic xenograft models of primary liver cancer, however, the tumours show less cell-to-cell variation in protein expression level and less overall protein expression relative to spontaneously-occurring HCCs.

Notably, mCherry-positive foci were detected in the lungs of 3 animals from this cohort, indicating spontaneous metastasis of the xenografted HuH-7 cells from the liver to the lung. eGFP fluorescence was not detected in these small lung metastases above the background fluorescence level of normal lung tissue (**Supplementary Figure 15**), suggesting that location of the tumour in the liver is necessary for mRNA-LNP uptake and expression.

### Delivery of mRNA-LNPs to secondary liver tumours

It is currently unknown whether cancer cells derived from other tissues can take up mRNA when engrafted in the liver. To model secondary liver cancer, we established derivatives of two human cancer cell lines, HCT116 (colorectal carcinoma) and A549 (lung adenocarcinoma), which constitutively express mCherry (**Supplementary Figure 11**). These cell lines were chosen because cancers of the colon and lung are among the most common primary sources for secondary liver tumours. Cells were engrafted into the livers of BALB/c nude mice by single direct injection (n=8-9), and after approximately 2 weeks, mice were injected with 10µg of eGFP mRNA-LNPs via the tail vein, then culled 24 hours later for tissue collection and imaging (**Figure 5a**).

**Figure 5:**
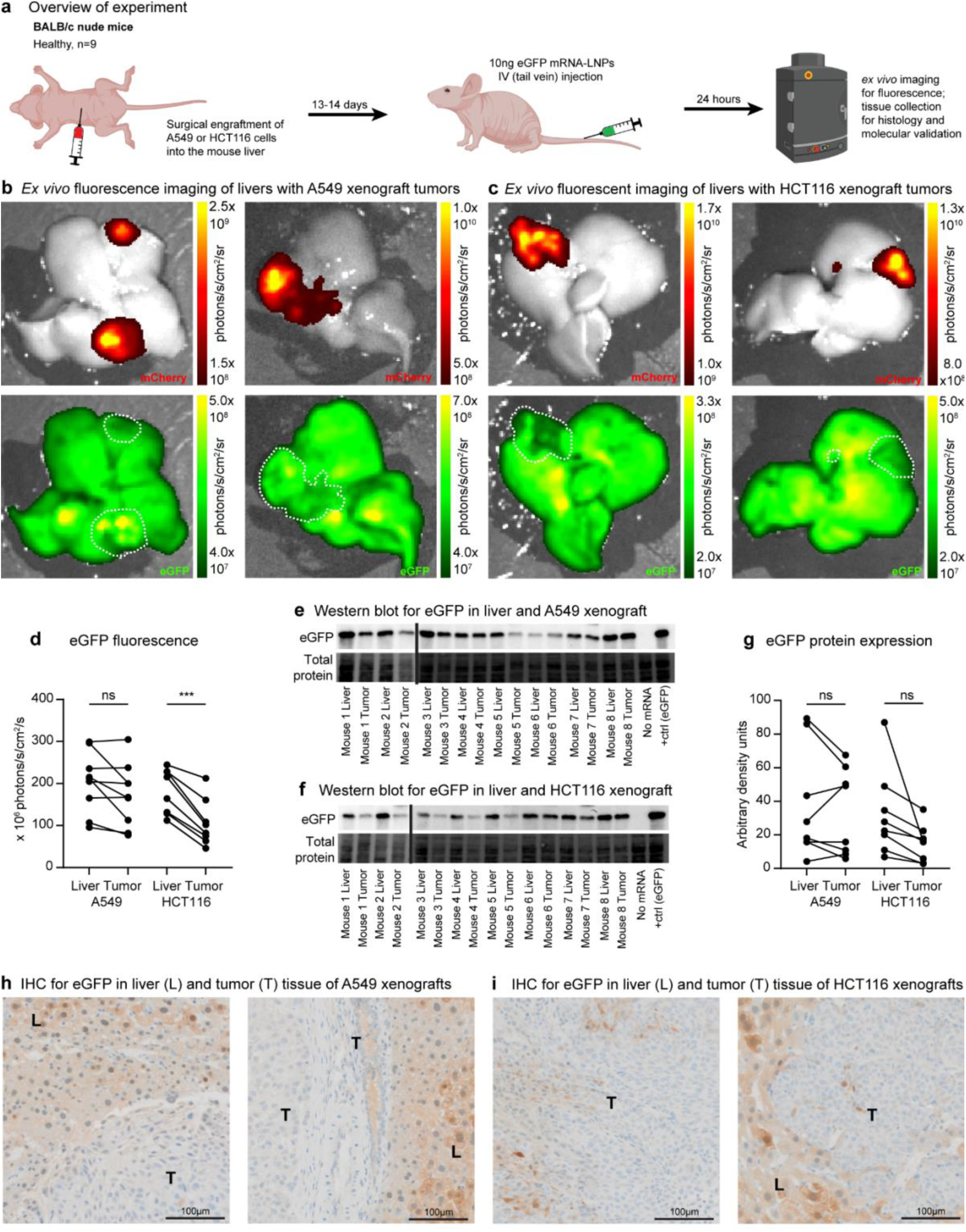
Delivery of mRNA-LNPs to secondary liver tumours. (**a**) We surgically engrafted A549 (human lung adenocarcinoma) and HCT116 (human colorectal carcinoma) cells, which constitutively express mCherry, into the livers of BALB/c nude mice. Tissues were analysed approximately 24 hours after an injection of 10µg eGFP mRNA-LNPs via the tail vein. **(b,c)** *Ex vivo* fluorescent imaging of entire livers with xenograft-derived tumours shows strong eGFP fluorescence in both liver tissue and tumours (marked by expression of mCherry). For A549-derived tumours **(b)**, fluorescence was visually comparable, while for HCT116-derived tumours **(c)**, fluorescent signal within the tumour was generally visibly lower than the adjacent liver tissue. **(d)** Quantitation of eGFP fluorescence for A549-derived tumours shows that fluorescence trends lower in tumours, but there is no significant difference (Paired t-test, n=9, t=2.269(8), p=0.0530). In HCT116-derived tumours, fluorescence is significantly reduced relative to liver tissue (Paired t-test, n=8, t=7.293(7), p=0.0002). **(e)** Western blotting detects eGFP from all livers and tumours of animals with A549-derived xenografts, with comparable or slightly lower band intensity in tumours relative to livers. **(f)** Likewise, Western blotting detects eGFP from all livers and tumours of animals with HCT116-derived xenografts, with slightly lower band intensity for most tumours relative to livers. **(g)** Quantitation of eGFP protein expression determined by Western blotting showed broadly comparable protein abundance in A549-derived tumours relative to liver tissue (Paired t-test, n=8, t=0.7250(7), p=0.4920). In HCT116-derived tumours, the abundance of eGFP protein trended lower but was not significantly different from liver tissue (Paired t-test, n=8, t=2.240(7), p=0.0601). **(h)** Immunohistochemistry images of liver and tumour tissue from two different animals with A549-derived tumours, showing faint staining for eGFP in tumour cells (T) and strong signal in adjacent liver (L). **(i)** Immunohistochemistry images for liver and tumour tissue from two animals with HCT116-derived tumours, showing faint to absent staining for eGFP in tumour cells (T) with strong signal in adjacent liver (L).

Using *ex vivo* fluorescence imaging, we again observed eGFP fluorescence well above background in the liver tissue of all mice. eGFP fluorescence was also detected in all tumours derived from both A549 and HCT116 (**Figure 5b,c**). We again used fluorescence imaging of tumour cross-sections to confirm mRNA expression within tumour interiors (**Supplementary Figure 16**). In A549 xenografts, fluorescence was comparable between liver (range 95.6-299.2 x 10^6^ photons/s/cm^2^/sr, average 203.4 x 10^6^ photons/s/cm^2^/sr) and tumour (range 77.1-304.9 x 10^6^ photons/s/cm^2^/sr, average 172.0 x 10^6^ photons/s/cm^2^/sr) (Paired t-test, n=9, p=0.0530) (**Figure 5b,d**). However, in HCT116 xenografts, fluorescence was significantly lower in the tumours (range 45.9-212.4 x 10^6^ photons/s/cm^2^/sr, average 106.2 x 10^6^ photons/s/cm^2^/sr) than in the liver tissue (range 112.1-244.5 x 10^6^ photons/s/cm^2^/sr, average 181.6 x 10^6^ photons/s/cm^2^/sr) (Paired t-test, n=8, p=0.0002) (**Figure 5c,d**). For both A549 and HCT116 xenografts, there was no correlation between tumour size and fluorescence intensity (**Supplementary Figure 17a,c**), but there was a moderate to strong correlation between the fluorescence intensity measured for the liver tissue and tumour from the same animal (**Supplementary Figure 17b,d**).

Western blotting confirmed the presence of eGFP protein in all tumours, again with considerable variability in band intensity (**Figure 5e-f**). There was no significant difference in eGFP protein abundance between liver tissue and tumours in the A549 group (range 4.2-89.3 and average 37.6 in liver, range 5.9-67.5 and average 33.6 in tumours) (Paired t-test, n=8, p=0.4920) (**Figure 5g**). Similarly, in the HCT116 group, there was no significant difference in eGFP abundance between liver and tumours, although the abundance in tumours trended lower (range 6.8-86.9 and average 32.3 in liver, range 2.9-35.1 and average 14.9 in tumours) (Paired t-test, n=8, p=0.0601) (**Figure 5g**).

Immunohistochemistry for eGFP in two A549 tumours (**Figure 5h, Supplementary Figure 18**) and two HCT116 tumours (**Figure 5i, Supplementary Figure 18**) showed weak expression of eGFP in most tumour cells and much stronger eGFP expression in adjacent healthy liver. These results indicate that mRNA-LNPs can be delivered to hepatic xenografts of non-liver cancer types. As with the HCC xenografts, these tumours show lower levels of eGFP protein expression relative to the liver tissue, and less cell-to-cell heterogeneity than was observed for spontaneous HCCs. mRNA-LNP uptake within secondary liver tumours substantially increases the potential scope of mRNA therapeutics for cancer.

## Discussion

In this study, we demonstrate the successful delivery of systemically-administered mRNA-LNPs to the diseased and cancerous mouse liver. We observed mRNA expression in healthy, fibrotic and cirrhotic mouse liver tissue, spontaneous hepatocellular carcinoma *in situ*, and xenograft models of HCC and secondary liver cancer. Using a custom machine-learning approach, we quantified the ubiquitous distribution of mRNA expression in hepatocytes in healthy liver, while showing that in the presence of fibrosis, cirrhosis and cancer, reporter expression varies both within and between samples. These findings demonstrate that intravenous administration of mRNA-LNP therapeutics can be a viable therapeutic strategy for liver disease and cancer, and suggest future directions for research to improve delivery of mRNA drugs.

Many nanoparticle-based carrier systems have been developed for the delivery of mRNA *in vivo*, with mRNA-LNPs being the most widely used. When mRNA-LNPs are injected intravenously, they rapidly adsorb the serum lipid transport protein ApoE^39^, which acts as a ligand for the low-density lipoprotein receptor (LDLR). This receptor is expressed on the surface of many cell types, but especially hepatocytes, which are the primary site for LDL clearance from the blood^40^. This feature of hepatocytes acts in conjunction with the fenestrated epithelium of the liver, which allows selective permeability of large particles into liver tissue which would be excluded from entry into other tissue types by the endothelial barrier^40^. Accumulation of nanoparticle drug carriers, particularly lipid-based carriers, in the liver is a well-documented phenomenon^40,41^, and many prior publications have shown that systemically-administered mRNA-LNPs are predominantly and abundantly expressed in the healthy liver, specifically by hepatocytes^26,27,42^. Delivery of mRNA to hepatocytes is a viable treatment option for several diseases, including genetic metabolic disorders where protein replacement occurring in hepatocytes, or secretion of the protein from hepatocytes into the bloodstream, is adequate to treat the disease. A notable example is mRNA-3927, a drug in clinical trials for treatment of propionic acidemia^31^.

To date, few studies involving mRNA-LNP delivery to the liver have considered the impact of hepatic fibrosis or cirrhosis. Liver fibrosis, characterized by the deposition of bands of extracellular matrix proteins within the liver tissue, results from chronic liver injury from a range of causes including chronic viral hepatitis, alcohol abuse, drug or chemical exposures, MASLD, and several genetic conditions. The end stage of this process is cirrhosis, characterized by extensive replacement of liver tissue with scar tissue so that liver function is significantly impaired^43^. In these disease states, the increased intracellular matrix within the space of Disse and capillarization of the sinusoids impedes blood flow and the transfer of molecules between sinusoids and hepatocytes. The applicability of mRNA therapeutics targeting the liver in the presence of chronic liver disease is thus partly dependent on the efficacy of mRNA-LNP delivery in the presence of these ultrastructural and morphological changes. One recent study described generally good mRNA-LNP delivery to the liver in mouse models of fibrosis and cirrhosis^32^. In this study, we build on these findings by using multiple orthogonal methods to describe the range of outcomes possible for mRNA-LNP delivery in the presence of severe chronic liver disease.

To investigate the impact of severe liver fibrosis or cirrhosis on mRNA-LNP delivery, we used the *Mdr2* knockout mouse, which develops progressive fibrosis and cirrhosis due to bile leakage^37,38^. Imaging of whole livers from *Mdr2*^−/−^ mice injected with eGFP mRNA-LNPs revealed significant eGFP fluorescence in the liver tissue, with both hypo– and hyperintense tumours evident. eGFP IHC performed on liver tissue sections from these animals showed considerable heterogeneity both within and between samples. Scattered hepatocytes with strong eGFP expression were a feature of virtually all sections, although their density varied considerably. Notably, we observed marginal to undetectable eGFP expression within fibrotic tissue tracts, which are comprised mainly of extracellular matrix and fibroblasts rather than hepatocytes. Our finding that the surface area of tissue sections with any reporter delivery was negatively correlated with the severity of fibrosis suggests that severe liver damage does impair delivery of mRNA-LNPs, however, moderately effective delivery was achieved even in mice with cirrhosis. Therefore, severe liver disease is unlikely to be a contraindication to the usage of mRNA drugs targeting the liver, although the dosage or frequency of administration may need to be adjusted to ensure adequate delivery in this patient population. Furthermore, mRNA therapies show promise for the direct treatment of liver injury and chronic liver disease. HNF4A mRNA inhibits the development of liver fibrosis in several animal models^32^, and expression of growth factors using mRNA-LNPs can reverse liver pathology and support engraftment of healthy hepatocytes to treat liver disease^27,44^.

Our study also systematically investigated mRNA-LNP delivery to liver cancer. *Mdr2*^−/−^ mice develop hepatocellular carcinomas with age, and we therefore examined reporter expression in spontaneous HCCs *in situ*. While a small number of reports previously described the use of mRNA-LNP based therapies to treat liver cancer in animal models^33–36^, the encoded therapeutic proteins act through non-cell-autonomous mechanisms of actions and delivery of mRNA-LNPs to tumour cells was unclear. We found that GFP was expressed in HCCs with a similar distribution and intensity to its expression in the liver tissue from the same animal. Cellular heterogeneity is a common feature of HCCs^45^, and this may explain our observation that several HCCs contained tracts of cells which did not express GFP. It would be valuable for future work to explore sequential delivery of multiple reporter mRNA-LNPs in order to determine whether repeated dosing of mRNA-LNPs would target the same or different cell populations. Interestingly, we found one steatotic HCC amongst the *Mdr2* KO tumours, and GFP expression was virtually absent from this tumour. However, robust GFP expression was observed in small steatotic regions within liver and tumour sections from other animals in this cohort. This finding warrants further investigation to determine whether steatotic livers or steatotic HCCs are especially poor targets for mRNA-LNP delivery.

As an alternative model for hepatocellular carcinoma, we also examined mRNA delivery in mice bearing xenografts of the human HCC cell line HuH-7. Fluorescence imaging showed comparable expression of eGFP in the livers and tumours of these animals. However, Western blotting showed reduced eGFP expression in tumours relative to adjacent liver tissue, including two tumours from which the eGFP band was barely detectable. Immunohistochemistry also showed much stronger eGFP staining in adjacent liver tissue than in tumours. We also considered the delivery of mRNA-LNPs to mice bearing xenografts of two human cell lines of non-liver origin, as a model for secondary liver cancer. Reporter expression was demonstrated in all tumours derived from both cell lines using multiple orthogonal methods, but expression was weak in comparison to the adjacent liver tissue.

Interestingly, delivery of mRNA-LNPs into the naturally-occurring HCCs of the *Mdr2* knockout mice was more efficient than delivery into any of the xenograft tumours. This is likely due to the difference in cell size and density; *Mdr2*^−/−^ tumours are comprised of much larger cells than xenografts derived from any of the three cell lines we considered. Studies of tissue penetration have found that nanoparticles larger than 20nm struggle to penetrate tumours due to their high cell density and interstitial fluid pressure^46–48^. High cell density and high tumour rigidity is typical of animal models bearing tumour xenografts derived from human cell lines, and the weaker delivery of mRNA-LNPs to these tumour models may represent an inherent limitation of the model system rather than the delivery technology. In contrast, the *Mdr2* knockout mouse model develops progressive liver disease which progresses through the stages of fibrosis, cirrhosis, and spontaneous development of HCCs; this is a physiologically relevant model which recapitulates the disease process which most commonly leads to HCCs in humans. Therefore, better performance of mRNA-LNPs in this model compared to xenograft models is a positive indicator of their applicability for cancer therapy. Our study also informs the choice of animal model for future research into mRNA-LNP therapeutics for liver cancer. The relatively weak delivery of mRNA-LNPs to orthotopic xenografts suggests that this model may underestimate the efficacy of mRNA cancer therapies, and that a mouse model which develops HCCs spontaneously is more likely to respond to treatment.

Nanoparticle composition for targeted mRNA delivery is an area of active research^49–51^. For instance, mRNA-LNPs with substituted lipid components enable selective targeting of the liver, lung and spleen^42^; other publications have described nanoparticle formulations which are optimized for RNA delivery to different types of liver cells^52^, or for co-delivery of siRNA, sgRNA and mRNA in a single LNP^33^. Excitingly, several technologies are in development which may improve the ratio of protein expression in tumour cells relative to liver cells, enabling therapeutic benefit with reduced mRNA-LNP dosage and reduced side effects related to expression of the mRNA in healthy hepatocytes. Cell-selective targeting including to extra-hepatic tumours can be achieved using bispecific antibodies which recognizes a tumour cell-surface marker and a component of the LNP^53^, and other studies have shown enhanced mRNA penetration into tumours when co-delivered with a therapeutic siRNA to reduce tumour rigidity^33^, as well as improved specificity of mRNA for tumours over hepatocytes by including binding sites for hepatocyte-specific miRNAs in the mRNA design^54^.

Another noteworthy finding of our study is the immune response, including PD-L1 elevation, in the liver 24 hours after injection of mRNA-LNPs. Exogenous mRNA is immunostimulatory through interactions with Toll-like receptors and cytosolic RNA sensors, and components of the mRNA-LNP are also immunostimulatory^50^. Our findings are consistent with previous reports of immune activation after mRNA-LNP administration. Importantly, one study reported that the inflammatory response in the liver was attenuated by reducing the dosage of mRNA-LNPs^32^. This highlights the importance of ongoing improvements to the design of mRNA medicines to enable reduction in mRNA dosage while still achieving therapeutic levels of protein expression.

In conclusion, we investigated reporter expression after systemic administration of mRNA-LNPs by the intravenous route results in mice. Building on previous findings showing strong expression of mRNA in hepatocytes, we have demonstrated mRNA expression in the healthy liver, fibrosis, cirrhosis, spontaneous hepatocellular carcinomas, and xenograft models of both primary and secondary liver cancer. Our results show that mRNA-LNP based cancer therapeutics are a promising drug modality for the treatment of liver cancer.

## Materials and methods

### Ethics statement

This study complies with the Australian Code for the Care and Use of Animals for Scientific Purposes. All procedures involving live animals were approved by the Institutional Animal Care and Use Committee of The University of Queensland (ethics approval certificates 2021/000492 and 2023/000234).

### Animals

Male BALB/cOzarc mice were obtained from Ozgene (Perth, Australia.)

Male BALB/c nude (BALB/c-*Foxn1^nu^*/Ozarc) mice were obtained from Ozgene (Perth, Australia.)

Male MDR2^−/−^ (FVB.129P2-*Abcb4^tm1Bor^*/J) mice were obtained from a breeding colony maintained by KRB and XL at the Pharmacy Australia Centre of Excellence (Brisbane, Australia.)

For imaging controls only, surplus or ex-breeder mice (male BALB/c, and male and female CD1) were obtained from a training colony maintained at the Australian Institute for Bioengineering and Nanotechnology (Brisbane, Australia.)

BALB/c and BALB/c nude mice used for baseline delivery and xenograft experiments were 8-10 weeks old when experiments commenced. *Mdr2*^−/−^ mice were 2-11 months old as indicated. BALB/c and CD1 imaging controls were 3-9 months old.

All animals used in this study were group housed (3-5 per cage) on a 12-hour light-dark cycle at ambient temperature, with *ad libitum* access to food and water and with enrichment items (cotton nesting material, shredded cardboard, carboard domes and popsicle sticks) provided in cages.

### Cells

HuH-7 human hepatocellular carcinoma cells were purchased from ATCC. HCT116 human colorectal carcinoma cells were a kind gift from Professor Michael McGuckin’s group. These cell lines were maintained in DMEM (Gibco 11965-092) supplemented with 10% fetal bovine serum (Gibco 10100147) and 1% penicillin/streptomycin (1,000U/mL, Gibco 15140122).

A549 human lung adenocarcinoma cells were a kind gift from the laboratory of Professor Helmut Schaider. A549 was maintained in RPMI 1640 medium (Sigma-Aldrich R8578) supplemented with 10% FBS, 1% pen/strep, 2mM L-glutamine (Thermo Fisher Scientific 25030081), 1mM sodium pyruvate (Thermo Fisher Scientific, 11360070), and 15mM HEPES (Thermo Fisher Scientific, 15630080).

All cells were maintained at 37°C with 5% CO_2_ in a humidified incubator. The cell lines used in this study are not listed in the International Cell Line Authentication Committee and National Center for Biotechnology Information Biosample database of misidentified cell lines. All cell lines used in this study tested negative for mycoplasma.

To generate derivatives of A549 and HCT116 which constitutively express mCherry, cells were seeded into 6-well plates and then co-transfected with pBRPB CAG-mCherry-IP plasmid (Addgene #106333) and the piggyBac transposase plasmid (PBase) at a molar ratio of 3:1 using Lipofectamine 3000 (Invitrogen L3000001) according to the manufacturer’s instructions. After 48 hours, puromycin selection was applied (6mg/mL for A549 and 5mg/mL for HCT116) and maintained for 2 weeks. To generate a derivative of HuH-7 which constitutively expresses mCherry and firefly luciferase, cells were seeded into 6-well plates and then co-transfected with pB-EF1a-FLuc-IRES-Puro plasmid and PBase at a 3:1 molar ratio using Lipofectamine 3000. After 48 hours, cells were selected with puromycin (5mg/mL) for 2 weeks. This cell line was then co-transfected with the pBRPB CAG-mCherry-IP plasmid and PBase at 3:1 molar ratio and selected with puromycin (5mg/mL) for a further 2 weeks. The BD FACSAria Fusion^TM^ instrument was used to select cells with red fluorescence, and this cell population was retained. All knock-in cell lines are routinely maintained in nonselective media.

To prepare cells for xenografting, cultures were grown to between 50-90% confluent and not passaged within the 24h prior to harvest. Cells were trypsinised and washed twice with PBS, then pelleted by gentle centrifugation and resuspended in undiluted Matrigel (Corning FAL354277). Cell suspensions were maintained on ice and used within 5 hours of preparation.

### Production of mRNA-LNP

mRNA-LNPs were produced according to a method published previously^55^. Briefly, a synthetic gene encoding eGFP was obtained from a custom DNA synthesis provider (Integrated DNA Technologies, Singapore). DNA was amplified using 2X Q5 PCR master mix (New England Biolabs M0494L) with a forward primer containing a T7 promoter, and a reverse primer encoding a 126-nt poly(A) tail. The PCR product was purified and used as a template for *in vitro* transcription of mRNA using T7 RNA polymerase (New England Biolabs M0251L), with 100% substitution of uridine residues by N1-methylpseudouridine (BOC Sciences 1429803-59-6.) The synthesised mRNA was purified, integrity verified by capillary electrophoresis, and function verified by transfection into mammalian cell culture. mRNA-LNPs were produced using SM-102 as the ionisable lipid (Moderna-like formulation) using the Nanoassemblr Ignite. mRNA-LNPs were buffer-exchanged into TBS by dialysis and concentrated by centrifugation on a size-exclusion column. A sample of concentrated mRNA-LNPs underwent quality control assessment using the Zetasizer to confirm that size, charge and polydispersity were within the expected range (**Supplementary Figure 19**). Remaining concentrated mRNA-LNPs were mixed with 10% sucrose as a cryopreservative and stored in single-use aliquots at –30C.

### mRNA delivery in healthy BALB/c mice

Male BALB/c mice 16 weeks old were injected via tail vein with 10µg of mRNA-LNPs diluted to a final volume of 100µL with TBS. Approximately 24 hours after injection, mice were euthanised and organs removed for imaging.

### mRNA delivery in *Mdr2*^−/−^ mice

4 groups of male *Mdr2*^−/−^ mice (n=4-5) were defined based on age: 2-3 months, 5 months, 8 months, and 11 months. Mice were injected via tail vein with 10µg of mRNA-LNPs diluted to a final volume of 100µL with TBS. Approximately 24 hours after injection, mice were euthanised and organs collected for histology and molecular analysis. A separate cohort of 11 month old *Mdr2*^−/−^ males (n=4) were used for imaging; these mice were injected via tail vein with 10µg of mRNA-LNPs diluted to 100µL with TBS, then euthanised approximately 14 hours later and organs removed for fluorescence imaging.

### mRNA delivery in an orthotopic xenograft model of human liver cancer

Male BALB/c nude mice (n=9) were provided with drinking water containing 200mg/L thioacetamide (TAA) for 8 weeks. 4-5 days after discontinuation of thioacetamide treatment, mice underwent surgery to engraft human cancer cells (Huh-7-FLuc-mCherry.) Briefly, mice were anaesthetised with isoflurane, then injected subcutaneously with 0.1mg/kg buprenorphine for analgesia and ophthalmic lubricant was applied. Mice were gently secured in dorsal recumbency using paper tape and the surgical site was prepared with 3 sequential povidone-iodine scrubs. A 2cm midline abdominal incision was made using scissors and the liver partially exteriorised onto sterile gauze swabs drenched in normal saline. 1×10^6^ cells suspended in 20µL of undiluted Matrigel (Corning FAL354277) were injected into the left lateral lobe of the liver using a 29g insulin syringe. Hemostasis was achieved using gentle pressure with cotton swabs wet with normal saline, then the abdomen was closed in 2 layers using 6/0 braided silk sutures (Ethicon 15232-ET) in a simple interrupted pattern. Subcutaneous buprenorphine 0.1mg/kg was provided twice daily for 3 days for postoperative pain relief.

Beginning 7 days postoperatively, mice underwent twice weekly bioluminescent imaging for monitoring of tumour size. Briefly, mice were injected subcutaneously with 100mg/kg D-luciferin (Vivo-Glo, Promega 1042), then imaged 15 minutes later using the IVIS Lumina X5 under light isoflurane anaesthesia. mRNA-LNPs encoding eGFP (10µg in 100µL of TBS) were injected via tail vein on each imaging day. Mice were euthanised 24 hours after mRNA-LNP injection when the estimated tumour size reached 1cc, or on day 15 post-xenograft. The 8 animals included in data analysis were therefore euthanised on day 12 (n=1) or day 15 (n=7). One animal required euthanasia for welfare criteria at day 7 (immediately after the first imaging session) and was therefore excluded from the study.

### mRNA delivery in xenograft models of secondary liver cancer

Male BALB/c nude mice were used for this experiment. No TAA treatment was performed. Mice underwent surgery for cancer cell engraftment according to the procedure described for the HuH-7 xenograft experiment. Mice received either 1×10^6^ A549-mCherry cells (n=9) or 2×10^6^ HCT116-mCherry cells (n=9). 13-14 days postoperatively, mRNA-LNPs encoding eGFP (10µg in 100µL of TBS) were injected via tail vein. Mice were euthanised approximately 24 hours later and organs removed for fluorescence imaging. One mouse in the HCT116 group required euthanasia for welfare criteria prior to mRNA injection and was therefore excluded from the study.

### Fluorescent imaging of mouse organs

Following euthanasia, organs were removed onto chilled metal trays covered with opaque black plastic. Fluorescent imaging was performed using the IVIS Lumina X5 operated using LivingImage 4.7.4 (Perkin-Elmer). Images were acquired using the following settings: Field of view B, fluorescence filter pair software presets for mCherry and eGFP, automatic exposure, cosmic ray correction on. For image display, the RedHot lookup table was used and intensity-to-colour scales were set with reference to uninjected controls to best capture the qualitative data contained within the images. For images representing eGFP fluorescence, a custom script was used to invert the R and G colour values of exported PNG images to create a ‘GreenHot’ image.

Data analysis was performed using LivingImage 4.7.4 (Perkin-Elmer). To obtain numerical data for fluorescence of organs and tissues, a region of interest (ROI) was drawn around each tissue or organ and the fluorescence (average radiant efficiency) was recorded. Background fluorescent signal (arising from tissue autofluorescence) was estimated for each organ by averaging the fluorescent signal obtained from that organ from 2-6 uninjected mice. For mice injected with eGFP mRNA-LNPs, eGFP fluorescence for each organ was determined by subtracting the average background fluorescence from the same organ.

### Tissue processing and immunohistochemistry

Tissue samples used for histology and immunohistochemistry were drop-fixed by placing them in 10% neutral buffered formalin (Sigma-Aldrich F5554-4L) as soon as practical after dissection and imaging. After 24 hours of fixation, tissues were transferred to 70% ethanol for short-term storage, then paraffin embedded using a standard 9-hour processing cycle. 4µm paraffin sections on plain glass slides were used for H&E and picrosirius red stains. 4µm paraffin sections on Uberfrost+ slides were used for IHC. Following EDTA-mediated antigen retrieval at pH 9, IHC to detect GFP was performed using the Ventana Discovery Ultra automated stainer (Roche Diagnostics) and rabbit anti-GFP primary antibody (Novus NB600-308) at 1:1000. Brightfield images were captured using the Olympus VS120 slide scanning microscope at 20X (H&E stain) or 40X (immunohistochemistry) magnification.

### Immunohistochemistry image analysis with a custom AI model, *StainDetectAI*

IHC images were analysed using *StainDetectAI,* which is a Convolutional Neural Network (CNN) machine learning model prepared using standard U-Net architecture^56^ with four encoding and decoding layers. Each layer included convolutional blocks with ReLU activation functions and batch normalisation. The model was implemented using the PyTorch 2.4 library^57^ and trained to distinguish stained and unstained areas of mouse liver and spontaneous HCC using annotated tiles extracted from GFP IHC slide images from the *Mdr2* KO dataset. ‘No staining’ was defined based on GFP IHC on liver sections from mice not injected with mRNA (**Supplementary Figures 1, 2**). Training was performed using the Adam optimiser^58^, a learning rate of 0.0001, the mean Intersection over Union (mIoU) loss function, and early stopping after 30 epochs of no improvement. The final model finished training with a cell classification accuracy of 94.4% and precision of 97.7%, and its predictions were manually validated.

To determine the stained surface area of mouse liver and tumour sections, microscopy images in .vsi format were extracted using a custom script and split into 512×512 pixel tiles. Tiles were analysed by the trained model and a prediction mask was generated. The number of pixels in each tile corresponding to a prediction of stained, unstained, and ‘other’ (e.g. empty sections of the slide) were then summed and recorded for each sample. The entire area of each section was considered, and where available, two or more consecutive sections from the same sample were averaged together to calculate the final staining percentage.

For the subsample analysis reported in **Supplementary Figure 5**, microscopy images in .vsi format were extracted using a custom script and split into 512×512 pixel tiles. Tiles were manually inspected and empty tiles, ‘edge tiles’ (containing 10% or more of the slide background at the edge of the tissue section), and tiles containing mostly fibrotic tissue or damaged regions of the section were deleted. For each liver, 10 of the remaining tiles were randomly selected, and regions of the image containing fibrotic bands were manually masked. *StainDetectAI* was run on the unmodified tile images, and then data corresponding to the masked regions was removed to calculate the stained surface area of the remaining parts of the tiles.

### Protein extraction and Western blotting

Tissue samples used for Western blotting were snap-frozen on dry ice as soon as practical after dissection and imaging. Frozen tissue samples (3-30mg) were disrupted in 200-300µL cold RIPA lysis buffer (Cell Signaling Technologies 9806) supplemented with HALT protease and phosphatase inhibitor cocktail (Thermo Fisher Scientific 78440), using a glass dounce homogeniser, then vortexed thoroughly and incubated on ice. Homogenates were centrifuged for 10 minutes at 10,000 x *g* and 4°C, then the supernatant was transferred to a new tube, vortexed, and aliquoted for storage at –80°C. Protein samples were quantified by BCA assay using the Pierce reducing agent compatible microplate BCA protein assay kit according to the manufacturer’s directions (Thermo Fisher Scientific 23252). SDS-PAGE was performed using 10µg of total protein per lane on Stainfree AnyKD mini TGX precast gels (Bio-Rad 4568126). Following activation of the Stainfree reagent using the ChemiDoc MP (Bio-Rad), protein was transferred onto low-fluorescence PVDF membranes (Bio-Rad) using the Trans-Blot Turbo (Bio-Rad) using the ‘1X Mini-TGX’ preset (2.5A and 25V for 3 minutes). Membranes were blocked in EveryBlot blocking buffer (Bio-Rad 12010020), then incubated overnight at 4°C with primary antibodies diluted in blocking buffer. Membranes were then washed, and probed using fluorescent secondary antibodies diluted in a mixture of 50% Tris-buffered saline and 50% EveryBlot blocking buffer with 0.02% SDS. Antibody details are listed in **Supplementary Table 2**. Blots were imaged using the ChemiDoc MP. Quantitative image analysis was performed using Image Lab 6.1 software and protein levels were normalised to total protein signal detected using the Stainfree reagent.

### RNA extraction, qRT-PCR and RNA sequencing

Tissue samples used for RNA extraction were snap-frozen on dry ice as soon as practical after dissection and imaging. Frozen tissue samples (3-30mg) were disrupted in 500µL room temperature Nucleozol (Macherey-Nagel 740404.200) using a glass dounce homogeniser. Water was added and samples centrifuged to remove contaminants according to the manufacturer’s directions. Supernatant was then mixed 1:1 with ethanol and RNA captured on a Zymo-Spin IC column (Zymo Research R1013). DNase treatment was performed on-column and RNA was washed and eluted according to the manufacturer’s directions. RNA integrity was verified using the Agilent 4200 TapeStation and samples used for sequencing were RIN 8 and above. Nonstranded sequencing libraries were prepared from the poly(A) fraction of the RNA samples using the VAHTS Universal V8 RNA-seq library prep kit for Illumina according to the manufacturer’s directions. Libraries were sequenced using the Illumina Novaseq 6000 platform as 2×150bp paired-end reads, at a depth of approximately 25 million reads per sample. Reads were aligned to the human genome (hg38) using STAR^59^, and gene counts were obtained via featureCounts^60^. Read counts were analyzed in Degust^61^ for differential gene expression using Voom/Limma, applying a threshold of ≥10 counts per million in at least three samples, FDR<1E-5 and fold change >2. Gene set enrichment analysis was performed using STRING^62^.

## Data availability

Mouse liver RNA-seq data will be uploaded to a public repository prior to publication. All other data supporting the findings presented in this paper are available from the corresponding author upon reasonable request.

## Code availability

Code and documentation created during this study is available on GitHub: https://github.com/SidHow/StainDetectAI

## Acknowledgements

The authors thank Dr Haotian Yang and Dr James Humphries for their generous advice and expertise. We acknowledge the facilities and scientific and technical assistance of the Centre for Advanced Imaging at the Australian Institute for Bioengineering and Nanotechnology. We gratefully acknowledge advice and technical services from the Histology Facility and Microscopy Facility at the Translational Research Institute, and UQ Biological Resources facilities. We acknowledge the following sources of funding and support: National Health and Medical Research Council (GNT2014002 and GNT1161832) to T.R.M., Australian Research Council (DE230100036) to S.W.C., Medical Research Future Fund (MRFCRI000063 and MRFCRI000089) to S.W.C. and T.R.M. National Collaborative Research Infrastructure Strategy (NCRIS), Therapeutic Innovation Australia (TIA) to T.R.M. and S.W.C., Tour de Cure to S.W.C., DC, L.J.L and T.R.M, and The University of Queensland to S.W.C. and T.R.M. The authors acknowledge the facilities and the scientific and technical assistance of BASE at The University of Queensland. BASE is supported by Therapeutic Innovation Australia (TIA). TIA is supported by the Australian Government through the National Collaborative Research Infrastructure Strategy (NCRIS) program.

## Author contributions

LJL and SWC conceived the study. LJL, KRB, XL and SWC designed experiments. LJL performed animal surgeries and imaging. YJG produced and validated modified cell lines. LJL, SUM, MV, YJG, NLC, CLDM, DKW and KRB performed tissue collection, sample processing and molecular experiments. GCM interpreted histology results. SAH designed and wrote software used for image analysis. XL and KRB manage the *Mdr2* knockout mouse colony and provided mice for the study. SWC, TRM, DAM and DHGC provided resources and supervised the study. LJL and SWC wrote the manuscript. All authors reviewed the manuscript.

---

**Supplementary Figure 1:**
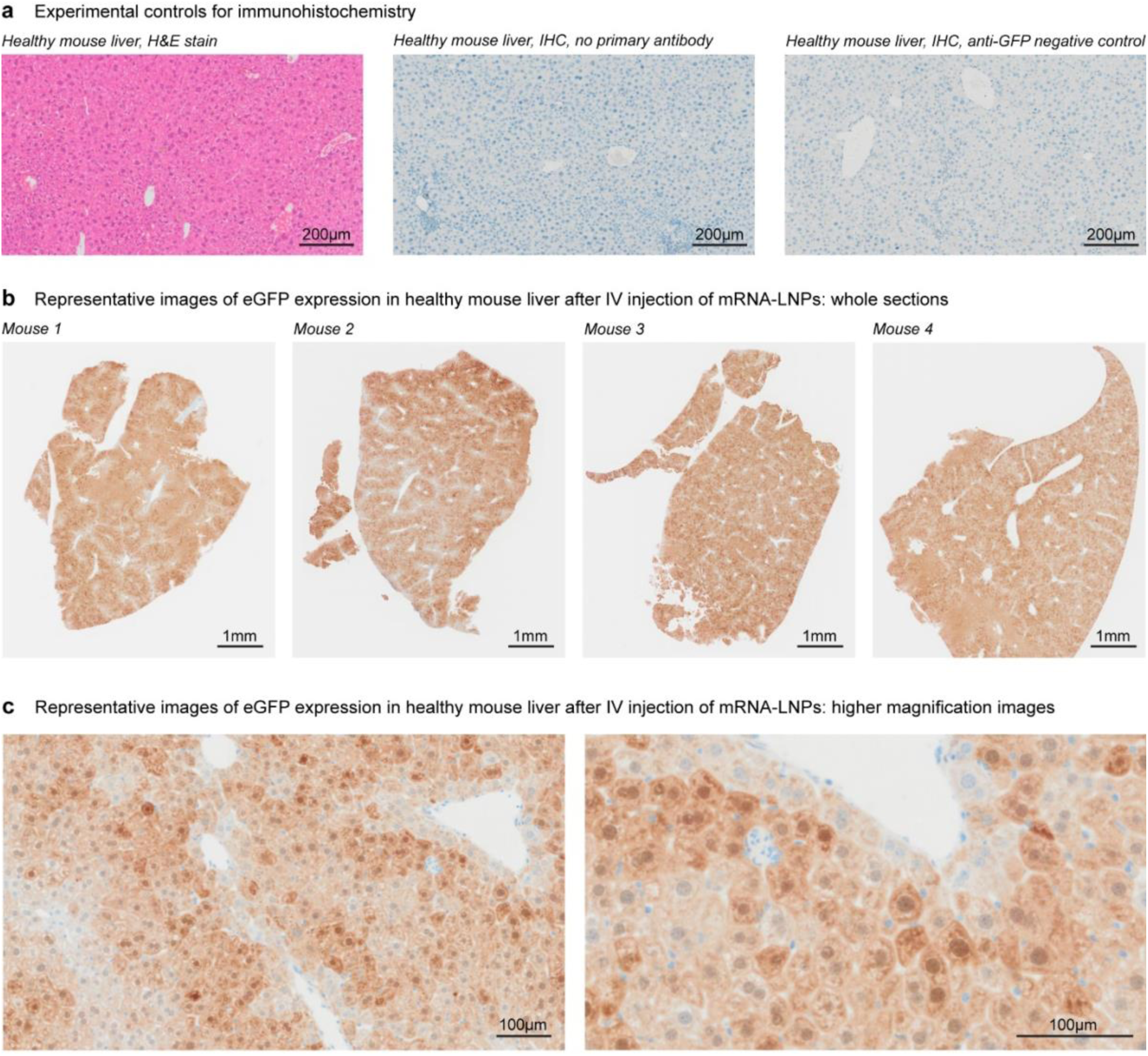
Additional histology images for healthy mouse liver. **(a)** H&E stain demonstrates normal morphology of healthy mouse liver. Immunohistochemistry controls demonstrate very low background of the assay. **(b)** whole section images taken from large pieces of mouse liver tissue (approx. 5 cubic mm) demonstrate that expression of eGFP from IV-injected mRNA-LNPs is strong and even throughout the liver tissue. **(c)** higher-magnification images demonstrate moderate to strong eGFP expression in all hepatocytes, with some expression visible in some endothelial cells.

**Supplementary Figure 2:**
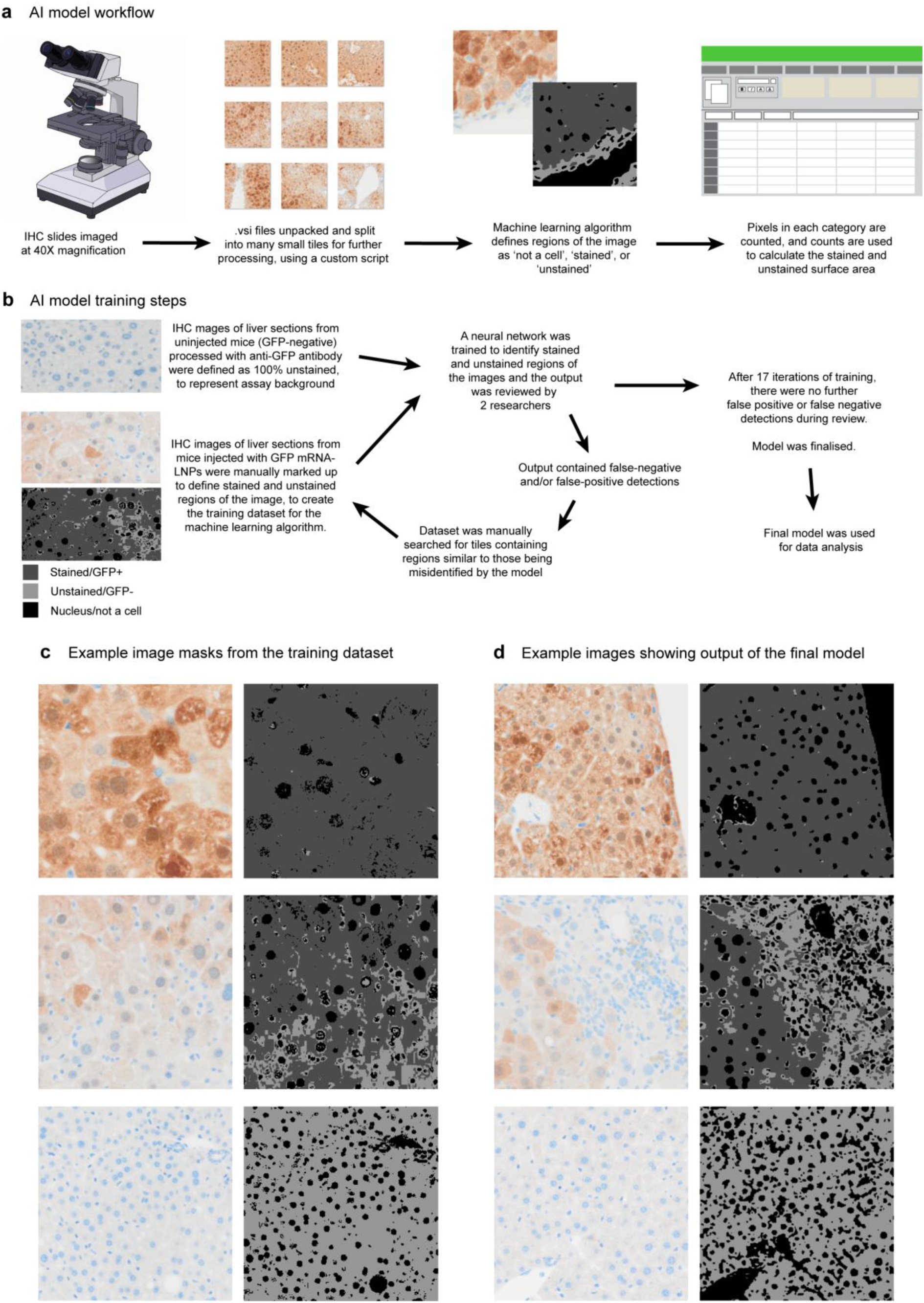
Overview of AI model used for IHC data analysis. **(a)** Flow chart of workflow for IHC data analysis using the custom AI model. **(b)** Flow chart of steps used to train the AI model to distinguish GFP-stained from unstained liver tissue. **(c)** Example image masks from the training dataset, used to provide input to the model. **(d)** Example output images from the final model, demonstrating its ability to accurately identify even light IHC staining with negligible false-positive detection.

**Supplementary Figure 3:**
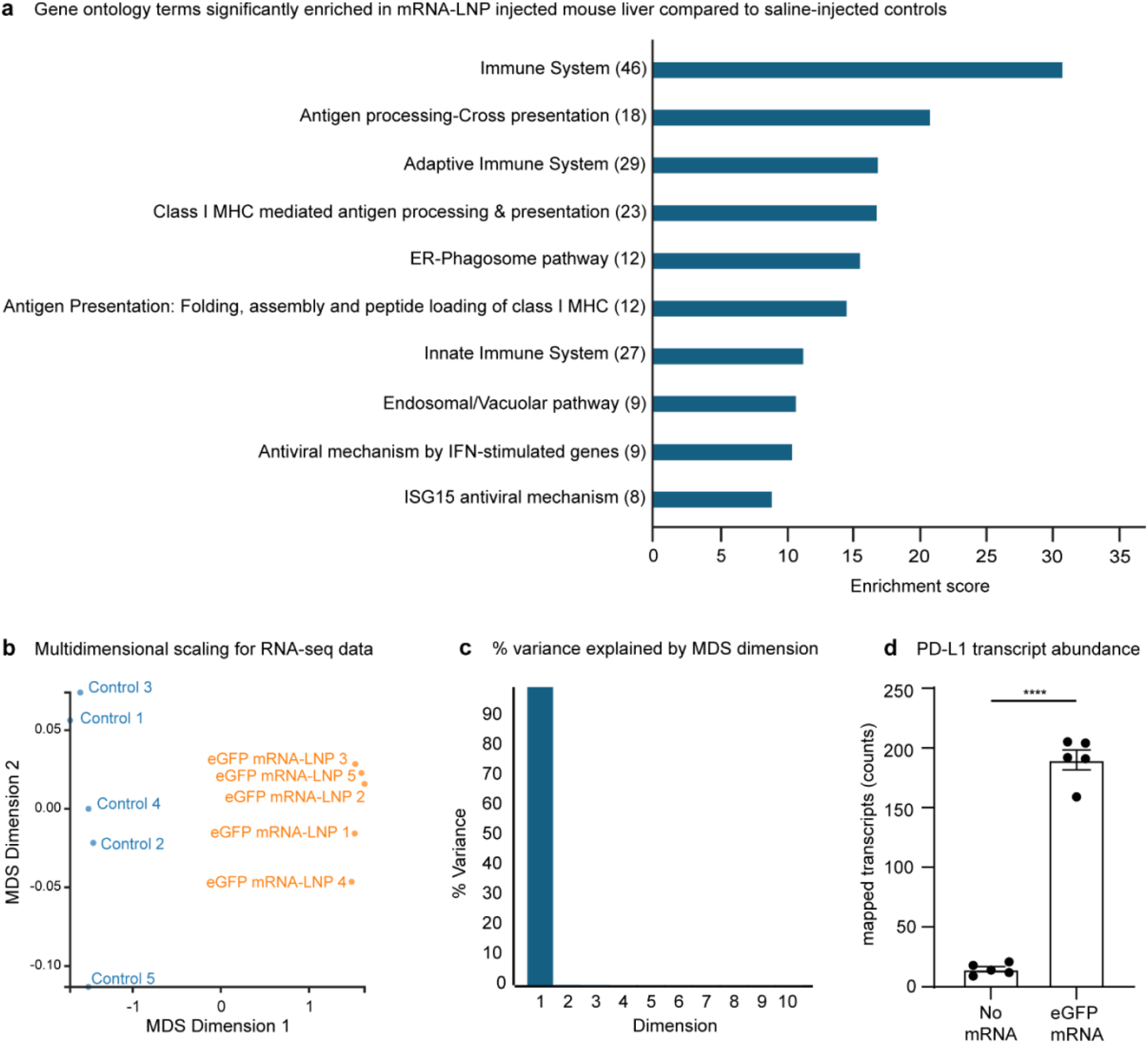
Additional results from RNA sequencing. **(a)** Gene ontology analysis of RNA sequencing data from the liver tissue of healthy mice injected with eGFP mRNA-LNPs compared to saline-injected controls. Enriched GO terms are related to the immune system and antigen processing and presentation. **(b)** Multidimensional scaling plot shows clear separation of mRNA-LNP injected animals from controls along MDS dimension 1. **(c)** MDS dimension 1 explains well over 90% of the variance in the data. **(d)** Abundance of CD274 (PD-L1) transcript was significantly increased in the mouse liver 24 hours after injection of eGFP mRNA-LNPs (Welch’s t-test, n=5, t=20.41(4.522), p<0.0001).

**Supplementary Figure 4:**
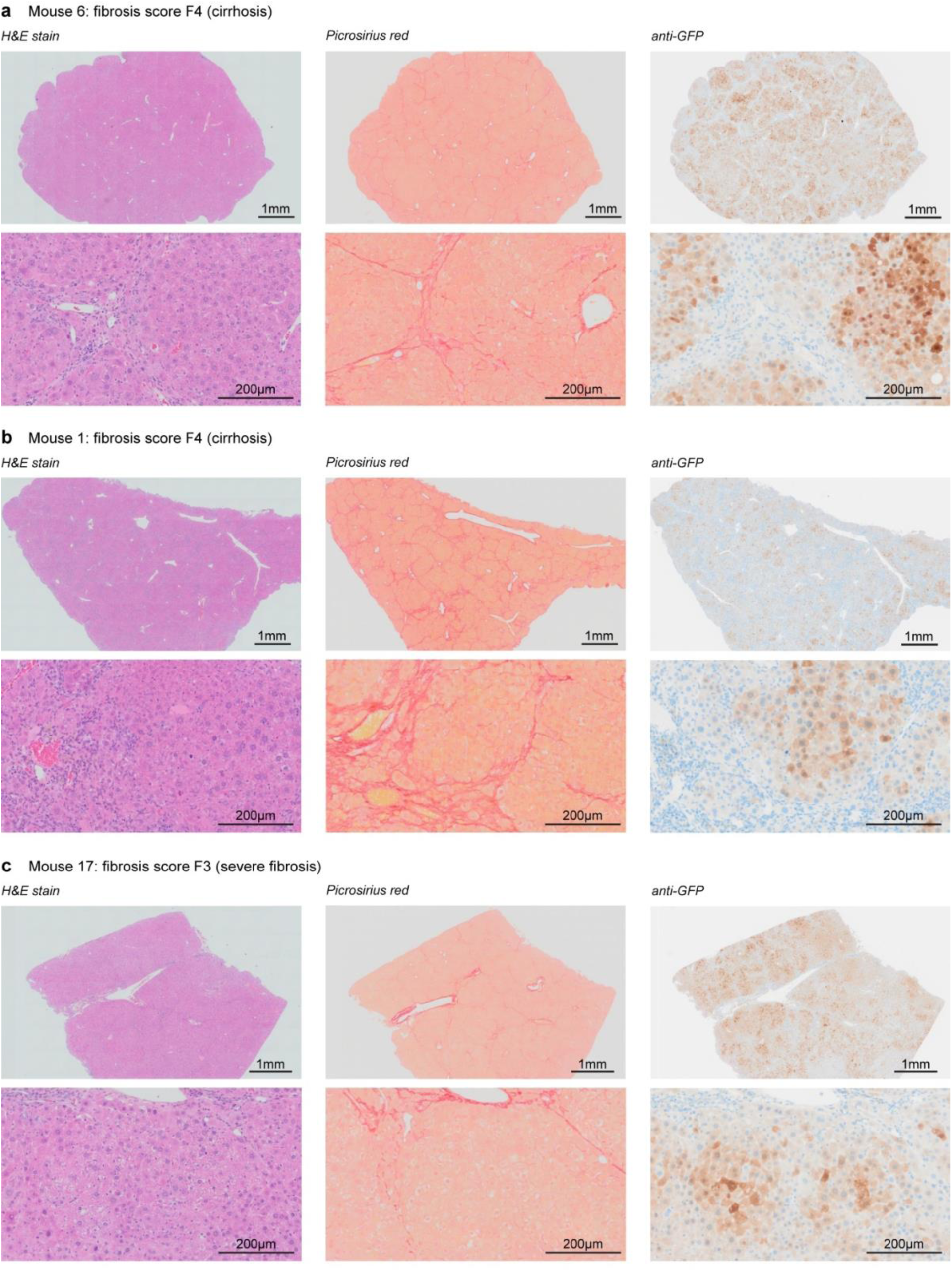
Additional histology images, Mdr2^−/−^ liver tissue. Mdr2^−/−^ mice develop bile leakage, severe fibrosis and cirrhosis. These images show the range of outcomes for liver disease severity and effectiveness of mRNA delivery. H&E stain (left) shows general tissue architecture, picrosirius red (centre) highlights collagen present in areas of fibrosis, and IHC for GFP (right) demonstrates delivery of mRNA-LNPs. **(a)** Typical delivery in an animal with cirrhosis (F4); 75% of the liver section surface area is positive for GFP. **(b)** Poor delivery in an animal with cirrhosis (F4); 44% of the liver surface area is GFP positive. **(c)** Typical delivery in an animal with severe fibrosis (F3); 84% of the liver surface area is GFP positive.

**Supplementary Figure 5:**
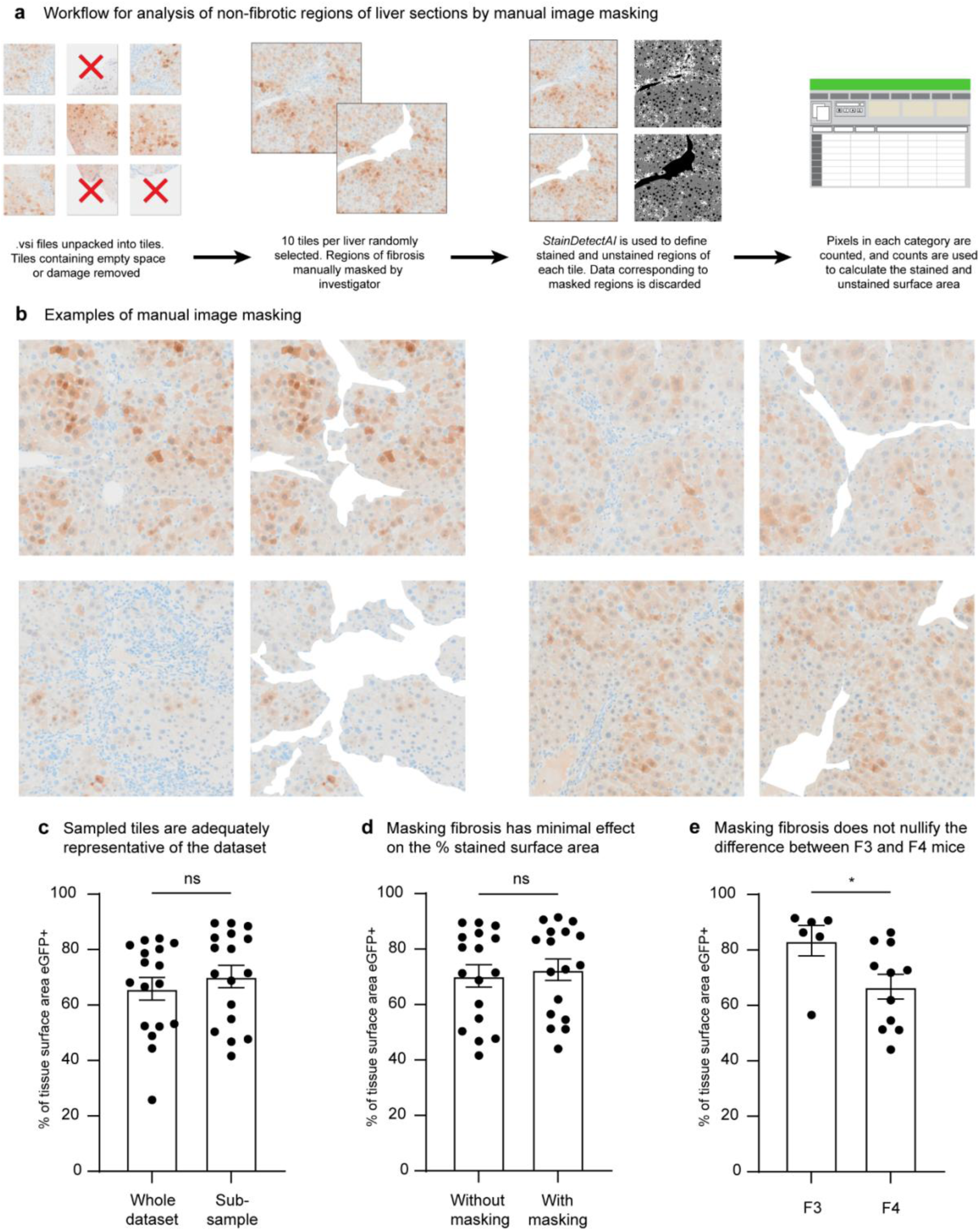
Analysis of IHC data subset with manual masking of fibrosis. **(a)** Flow chart of workflow for analysis of a random sample of IHC data from Mdr2^−/−^ mice with manual masking applied to the images, to consider the eGFP expression distribution in only non-fibrotic regions of the tissue sections. **(b)** Example of IHC image tiles without (left) and with (right) the masking applied. **(c)** Comparison of stained surface area (measured using *StainDetectAI*) for the random dataset sample in comparison with the whole dataset indicates that the random sample is representative of the dataset as a whole (Mann-Whitney test, n=17, U=115, p=0.3180). **(d)** Comparison of stained surface area for the random dataset sample with and without masking shows that there was no significant difference in the percentage of staining identified after masking fibrotic regions (Mann-Whitney test, n=17, U=124, p=0.4901). **(e)** Consistent with the findings of the whole-dataset analysis, in the masked sample there is a significant reduction in stained surface area for liver sections from animals with a fibrosis score of F4, relative to those scoring F3 (Mann-Whitney test, n=17, U=8, p=0.0103).

**Supplementary Figure 6:**
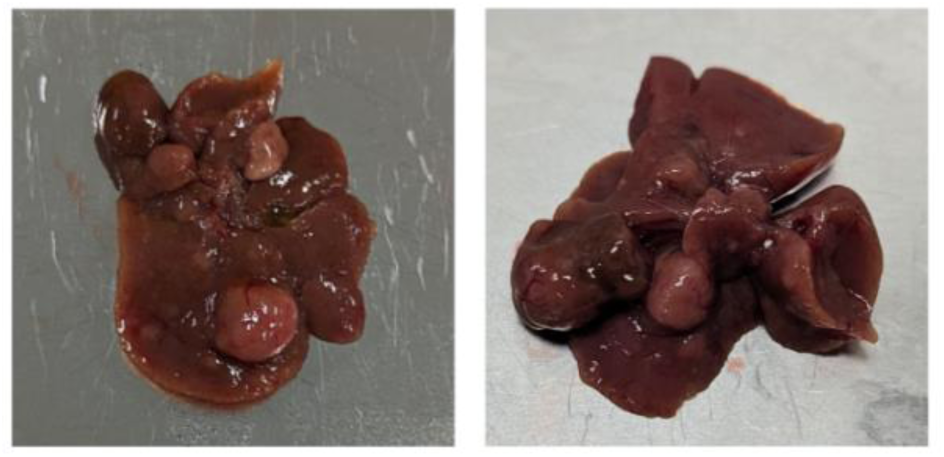
Photographs of Mdr2^−/−^ mouse livers. Representative photographs of whole livers from 11 month old male Mdr2^−/−^ mice, showing multiple large hepatocellular carcinomas visible on external surfaces of the liver.

**Supplementary Figure 7:**
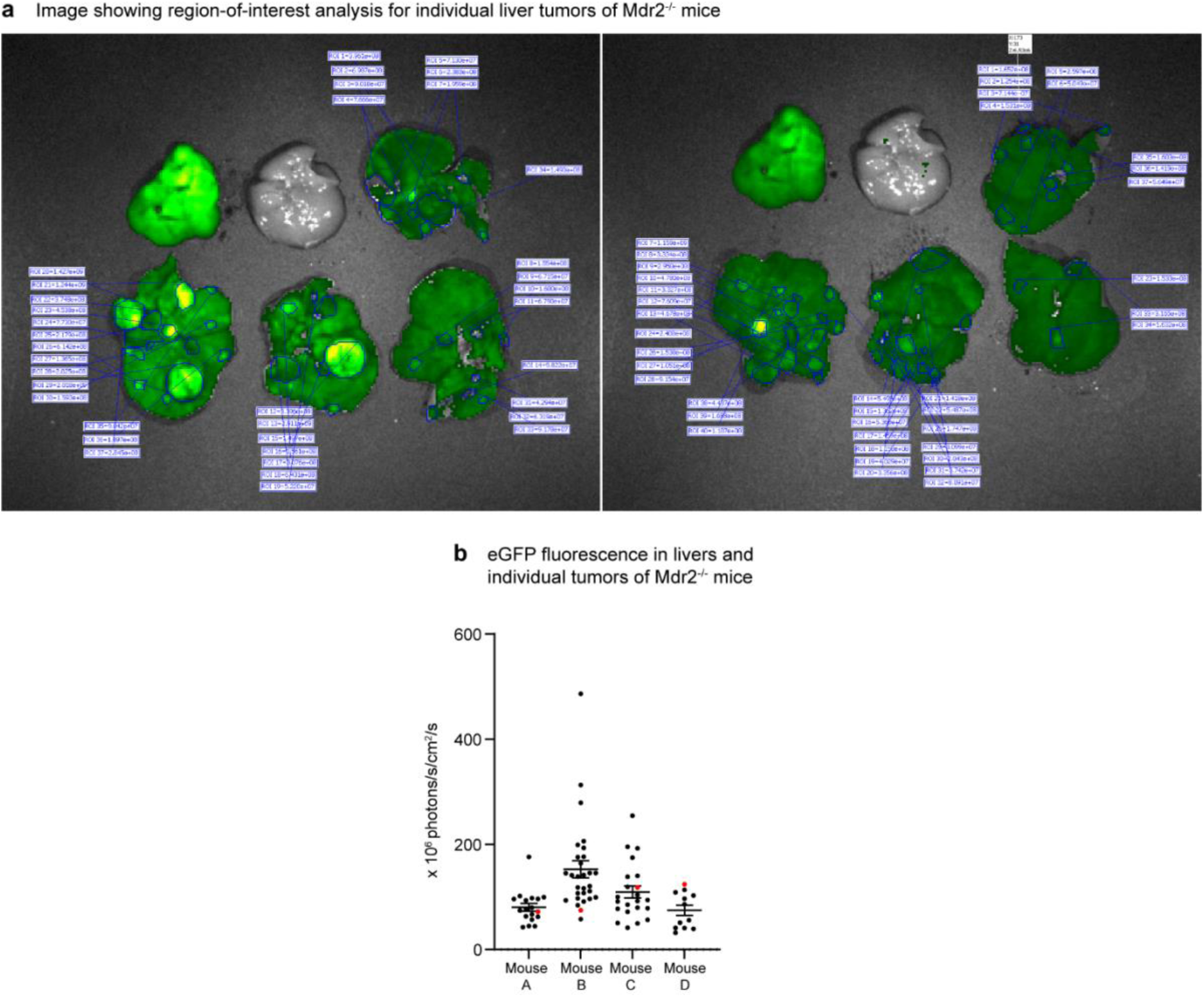
Region-of-interest analysis of individual tumors of Mdr2^−/−^ mice. **(a)** To analyse the fluorescence of individual liver tumors from Mdr2^−/−^ mice, regions of interest were drawn around identifiable tumors on both sides of the livers using the LivingImage software package. **(b)** Fluorescence of each region was calculated for each of four mice. The red point in each column represents the average fluorescence of the entire liver, including all tumors.

**Supplementary Figure 8:**
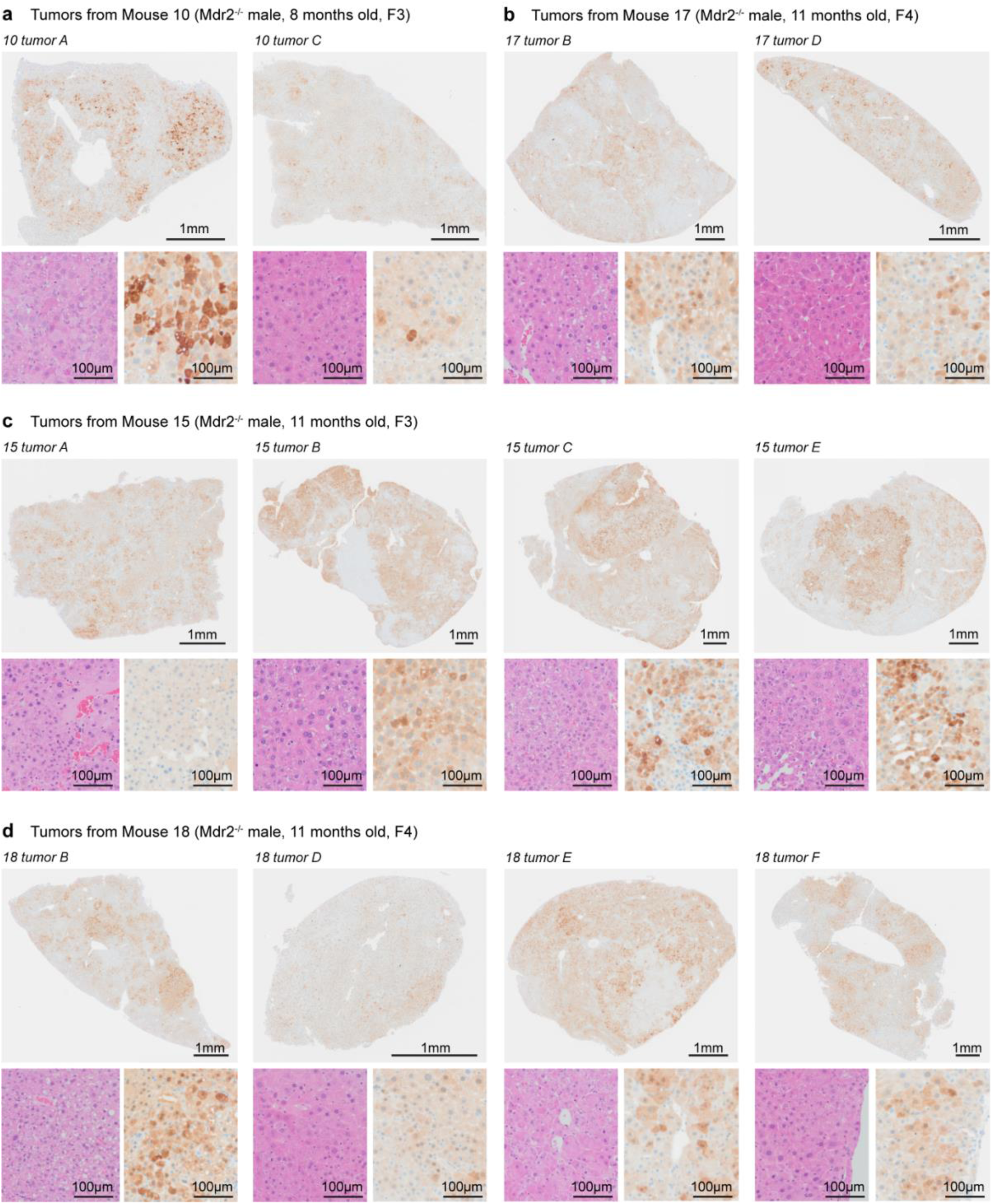
Additional histology images, Mdr2^−/−^ HCCs. Mdr2^−/−^ mice develop numerous HCCs with age. These images show the range of outcomes for mRNA-LNP delivery to various HCCs from four different male Mdr2^−/−^ mice.

**Supplementary Figure 9:**
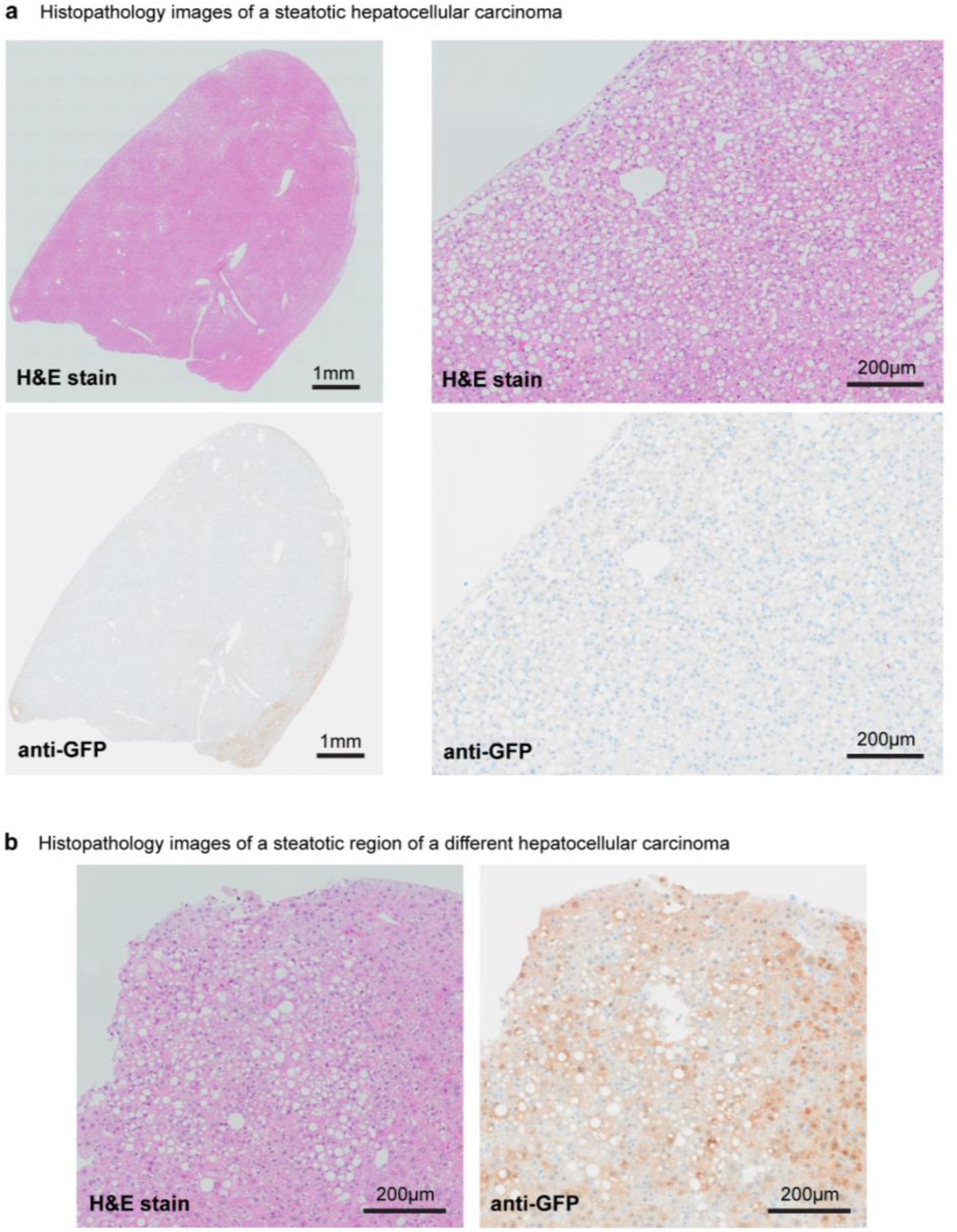
Delivery of mRNA-LNPs to steatotic HCC. **(a)** mRNA-LNP delivery to a steatotic HCC from an 11 month old male Mdr2^−/−^ mouse was notably poor, with eGFP expression only observed in a narrow band around the edge of the tumor. **(b)** Presence of steatosis is compatible with good mRNA-LNP delivery and expression, as demonstrated by strong eGFP expression in a steatotic region of a different HCC.

**Supplementary Figure 10:**
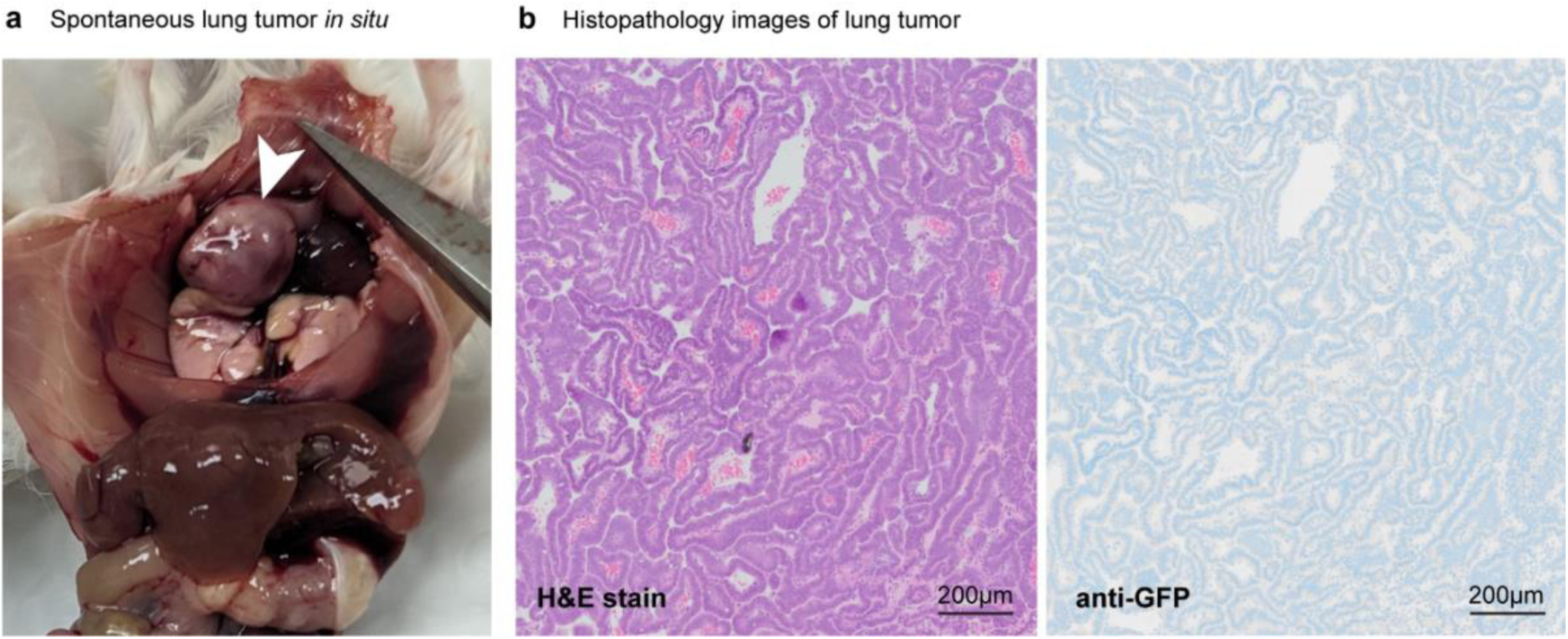
mRNA-LNPs were not delivered to one lung adenocarcinoma. **(a)** A large, spontaneous adenocarcinoma of the lung was recovered from an 11 month old male Mdr2^−/−^ mouse. **(b)** Delivery of mRNA-LNPs to this tumor was negligible, with eGFP staining not detectable above background.

**Supplementary Figure 11:**
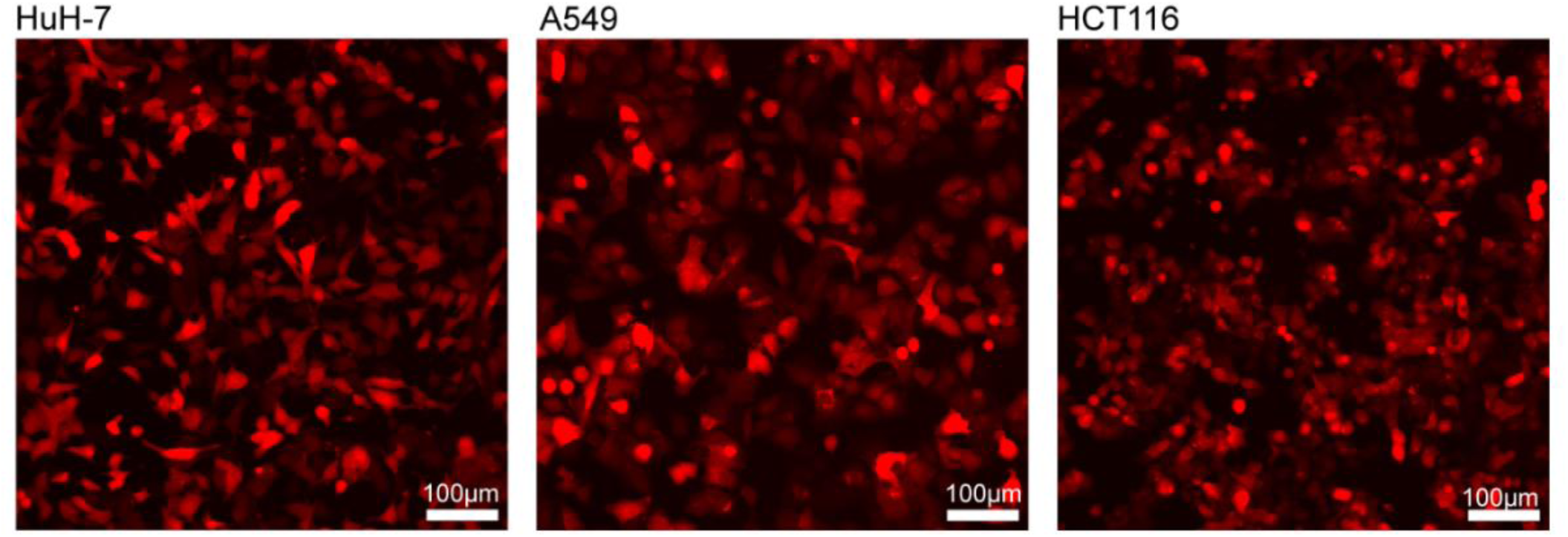
Demonstration of mCherry expression in cell lines. Fluorescence images showing strong expression of mCherry in knock-in cell lines used for animal experiments.

**Supplementary Figure 12:**
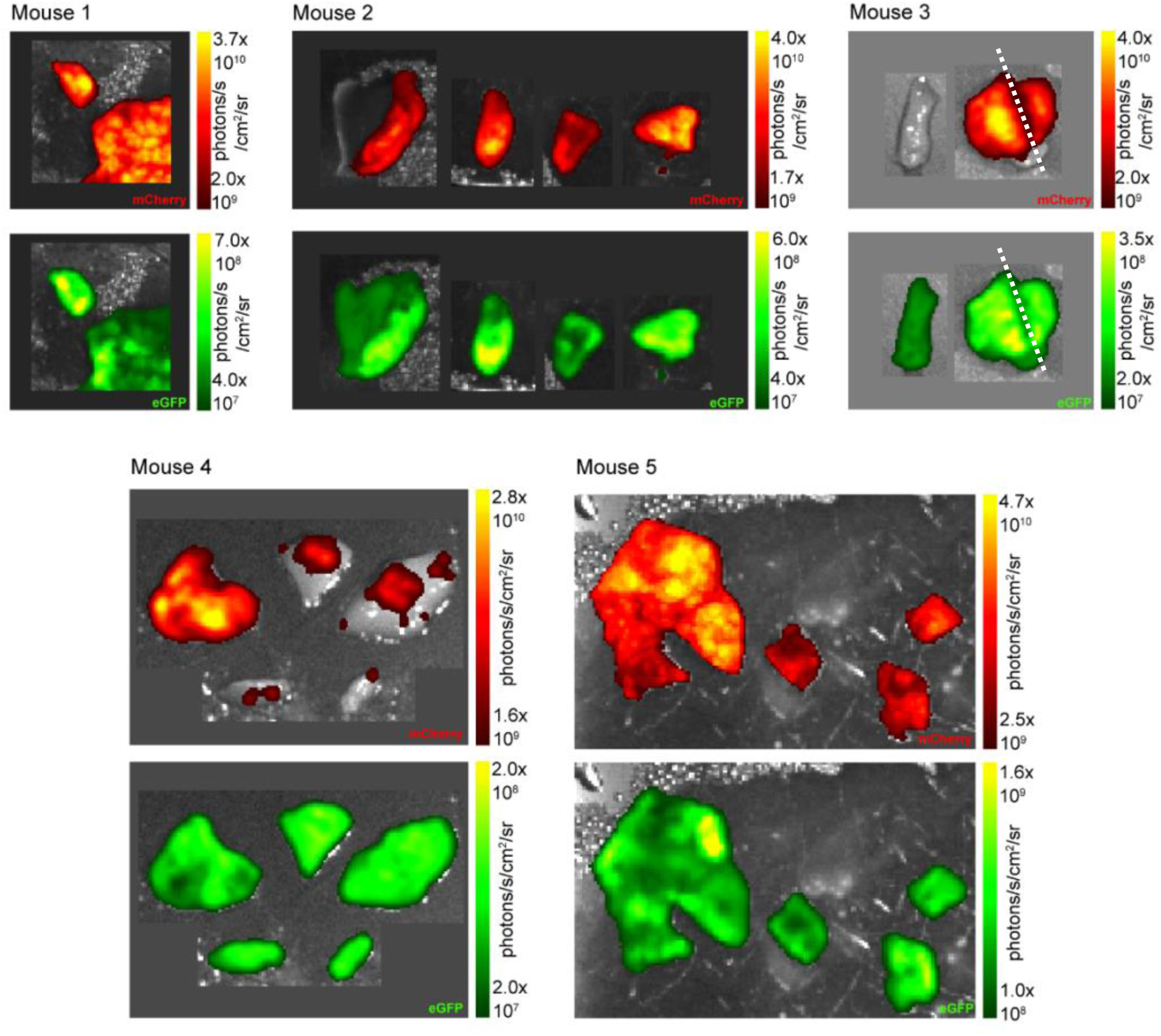
Fluorescence images for internal regions of liver tumors derived from xenografted HuH-7 cells. *Ex vivo* fluorescence imaging of liver tumors derived from HuH-7 human hepatocellular carcinoma cells xenografted into BALB/c nude mice. During tissue collection and imaging, tumors were cut into multiple pieces to provide samples for multiple analytical techniques, and images were taken to verify tissue identity and evenness of eGFP expression in internal regions of the tumors. Red (mCherry) identifies tumors, green identifies eGFP. Images show small sections of HuH-7 derived tumors with cut surfaces facing the camera. In most examples, even fluorescent signal is observed across the cut edge of the tumor, illustrating that eGFP is present throughout the tumor and not merely at the margins or in overlying healthy liver tissue.

**Supplementary Figure 13:**
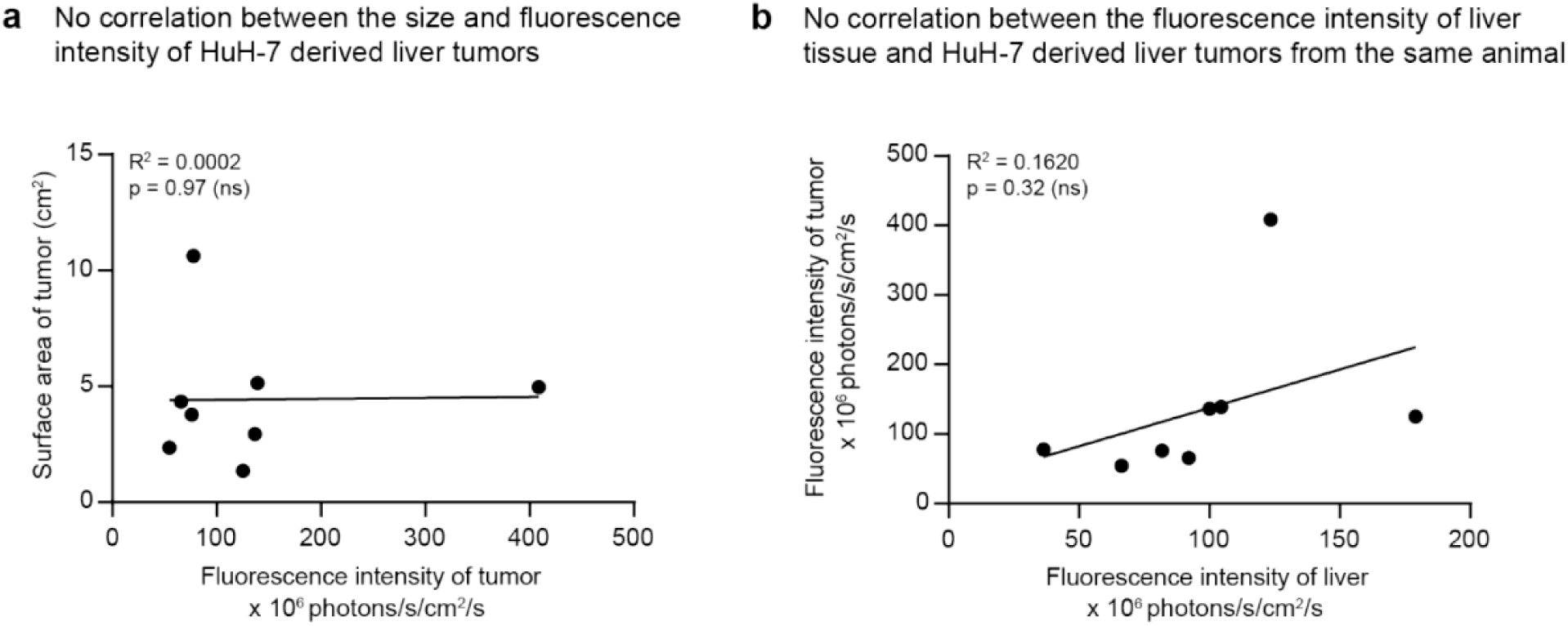
Additional analysis of fluorescence imaging data for HuH-7 derived xenografts. **(a)** There is no correlation between the size of HuH-7 derived liver tumors and their fluorescence intensity (Simple linear regression, slope not significantly different from zero (p=0.97)). **(b)** In mice bearing HuH-7 liver xenografts, there is no correlation between the fluorescence intensity of the liver tissue and the HuH-7 derived tumor (Simple linear regression, slope not significantly different from zero (p=0.1620)).

**Supplementary Figure 14:**
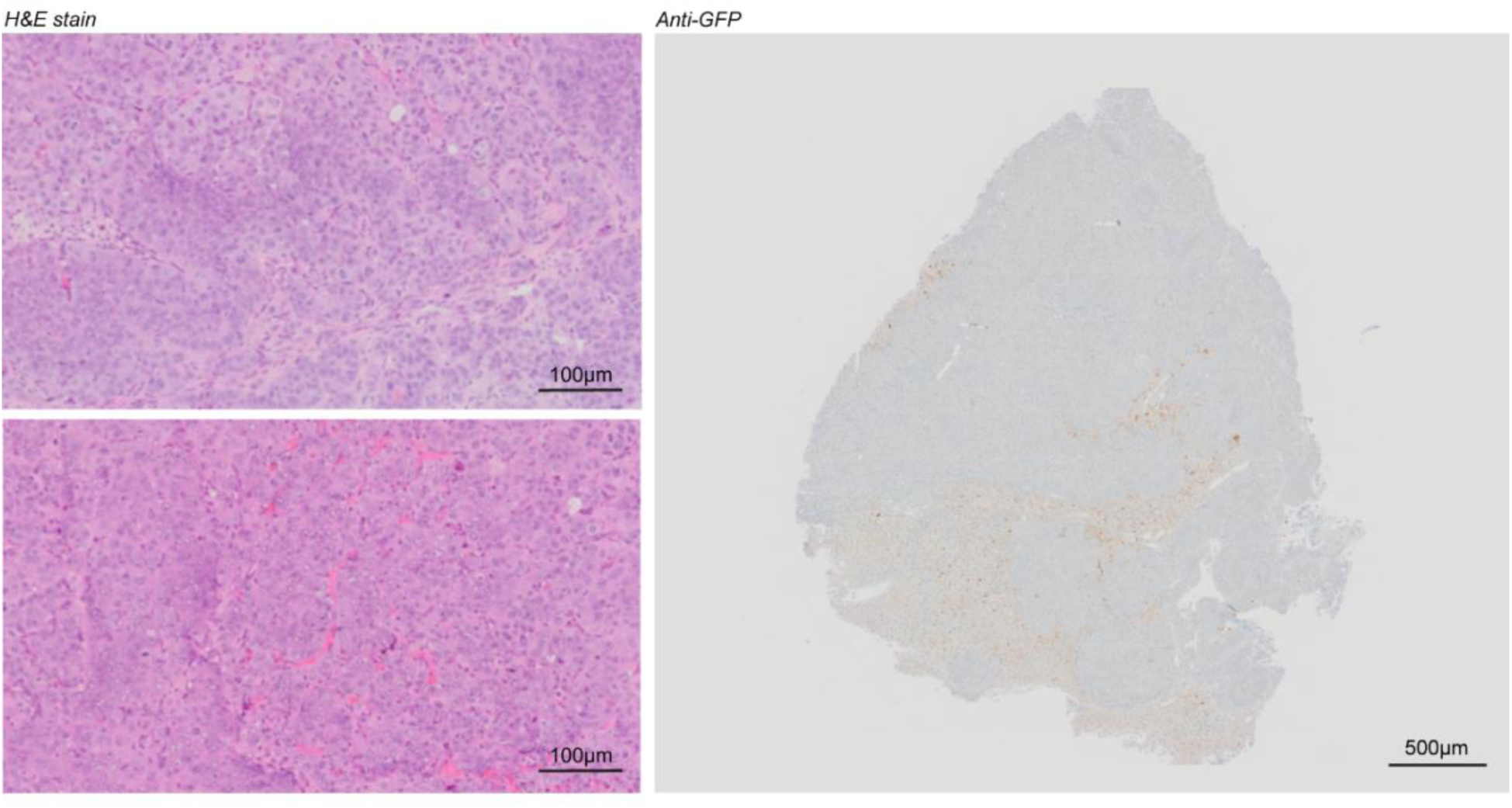
Additional histology images for HuH-7 derived xenograft liver tumor. H&E stain from representative regions of the tumor demonstrates tissue architecture. Anti-GFP stain shows low-level expression of eGFP throughout the tumor, indicating successful delivery of mRNA-LNPs. Note the presence of residual hepatocytes/liver tissue at the lower margin of the tumor. Smaller cells and higher cell density within the xenograft relative to the healthy liver makes the tumor tissue appear more blue; the colour of the cytoplasm (indicating eGFP staining) is comparable.

**Supplementary Figure 15:**
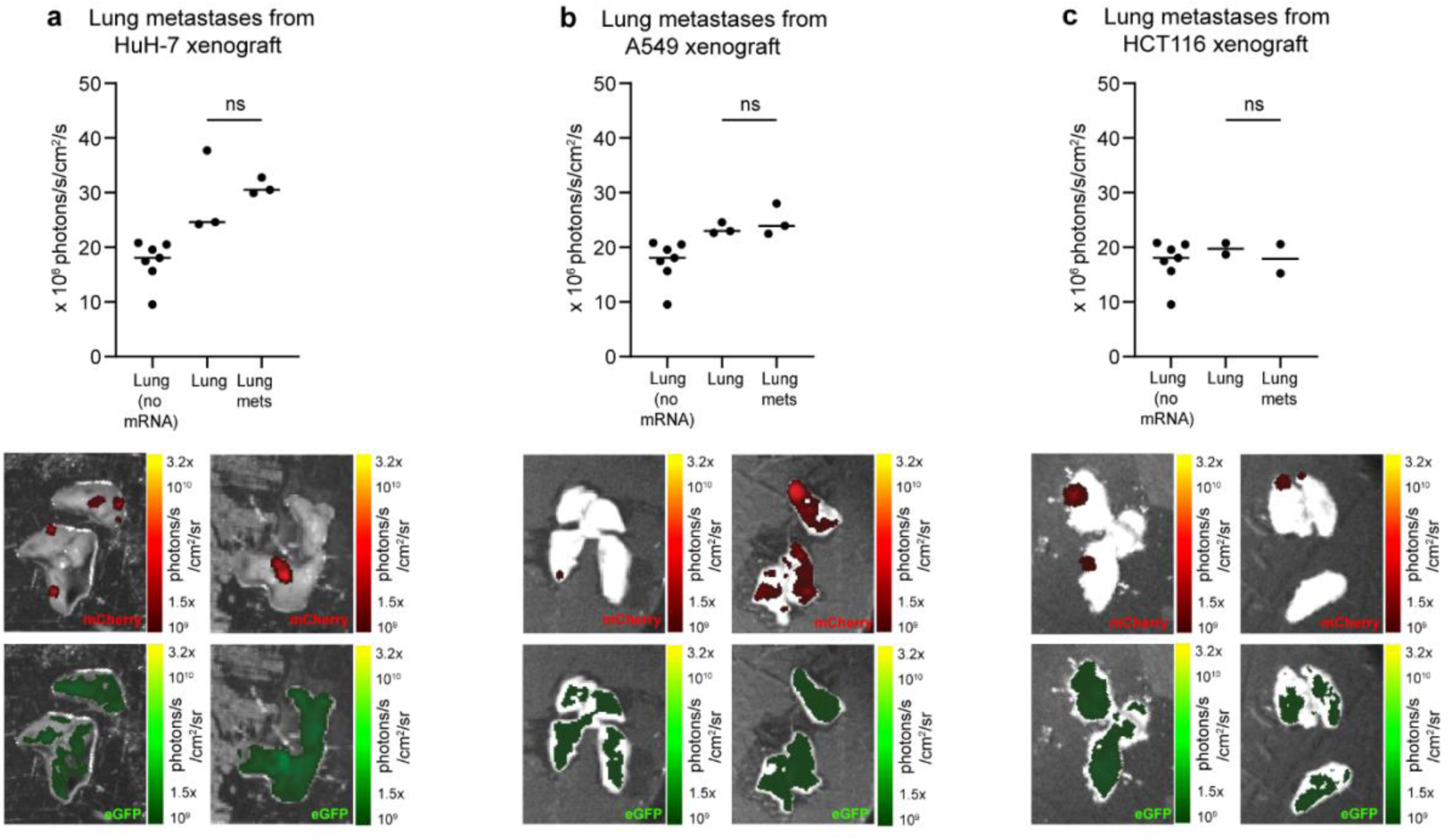
Fluorescence analysis of spontaneous lung metastases from mice bearing liver xenografts. In some animals bearing liver xenografts of human cancer cell lines, small metastases to the lungs were present (marked by red fluorescence). Region-of-interest analysis was used to compare the fluorescence of the metastases to the background fluorescence of the lung. There was no significant difference in fluorescence for any cell line, indicating that there is unlikely to be any substantial delivery of mRNA-LNPs to tumors of exogenous origin present in the lung. **(a)** No difference in fluorescence for HuH-7 derived lung metastases (Wilcoxon matched-pairs test, n=3, W=2, p=0.7500). **(b)** No difference in fluorescence for A549-derived lung metastases (Wilcoxon matched-pairs test, n=3, W=2, p=0.7500). **(c)** No difference in fluorescence for HCT116-derived lung metastases (Mann-Whitney test, n=2, U=1, p=0.6667).

**Supplementary Figure 16:**
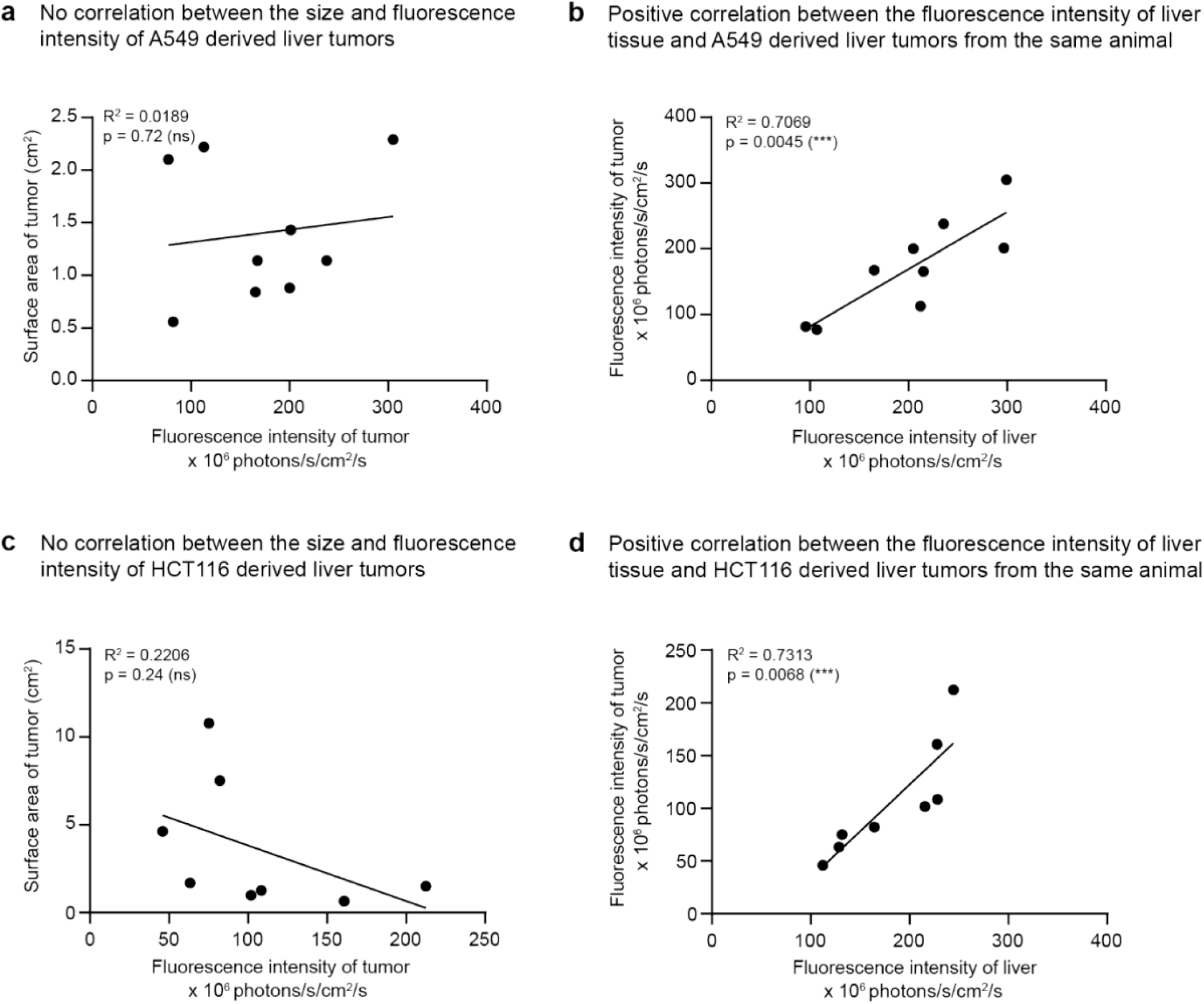
Additional analysis of fluorescence imaging data for A549 and HCT116 derived xenografts. **(a)** There is no correlation between the size of A549 derived liver tumors and their fluorescence intensity (Simple linear regression, slope not significantly different from zero (p=0.72)). **(b)** The fluorescence intensity of liver tissue and A549 tumor tissue in the same animal is positively correlated (Simple linear regression, slope significantly different from zero (p=0.0045), 70.69% of the variance is explained by the linear model (R^2^=0.7069)). **(c)** There is no correlation between the size of HCT116 derived liver tumors and their fluorescence intensity (Simple linear regression, slope not significantly different from zero (p=0.24)). **(d)** The fluorescence intensity of liver tissue and HCT116 tumor tissue in the same animal is positively correlated (Simple linear regression, slope significantly different from zero (p=0.0068), 73.13% of the variance is explained by the linear model (R^2^=0.7313)).

**Supplementary Figure 17:**
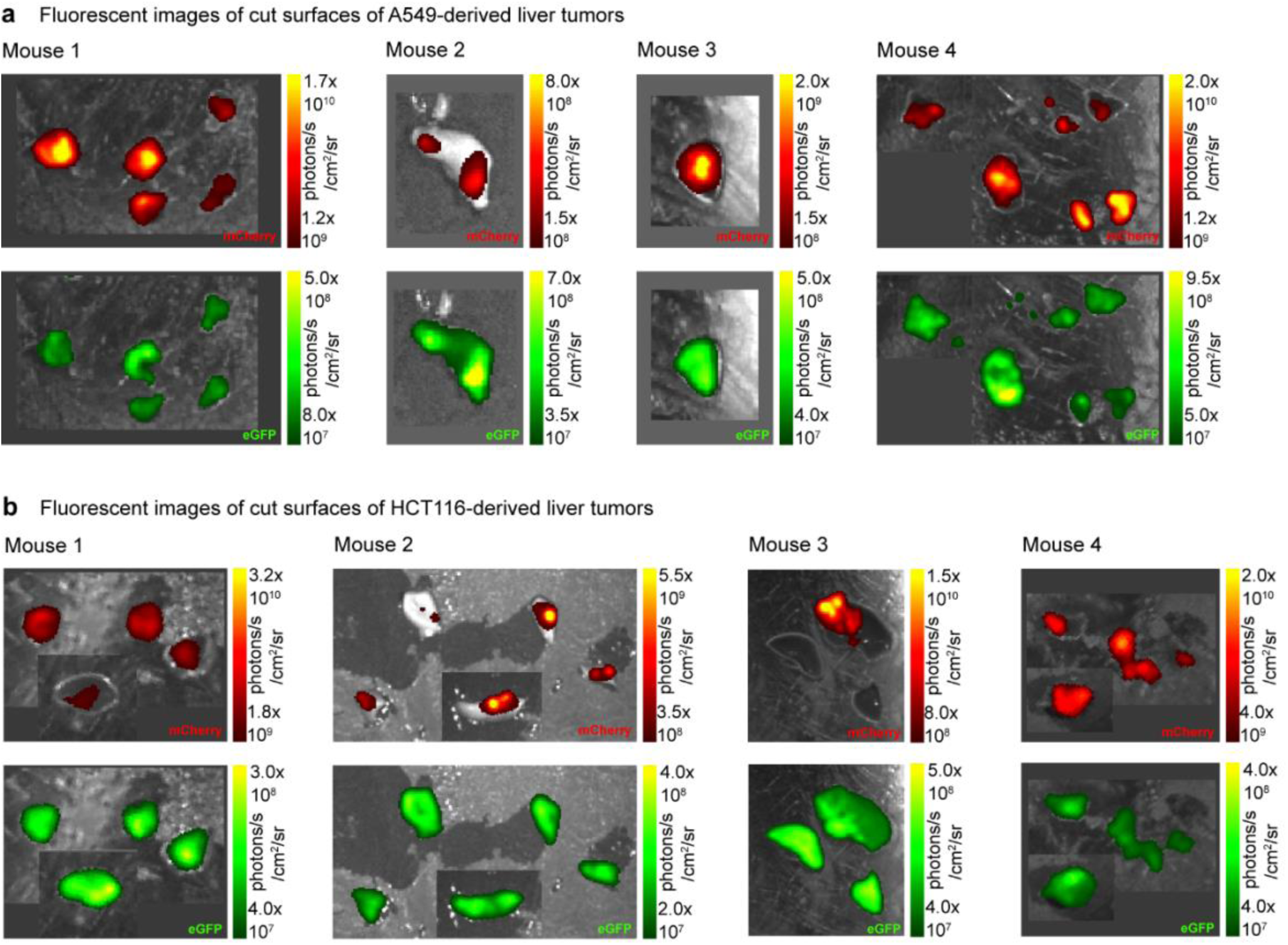
Fluorescence images for internal regions of liver tumors derived from A549 and HCT116 xenografts. *Ex vivo* fluorescence imaging of liver tumors derived from A549 **(a)** and HCT116 **(b)** cells xenografted into the livers of BALB/c nude mice. During tissue collection and imaging, tumors were cut into multiple pieces to provide samples for multiple analytical techniques, and images were taken to verify tissue identity and evenness of eGFP expression in internal regions of the tumors. Images show small sections of with cut surfaces facing the camera. In most examples, even fluorescent signal is observed across the cut edge of the tumor, illustrating that eGFP is present throughout the tumor and not merely at the margins or in overlying healthy liver tissue.

**Supplementary Figure 18:**
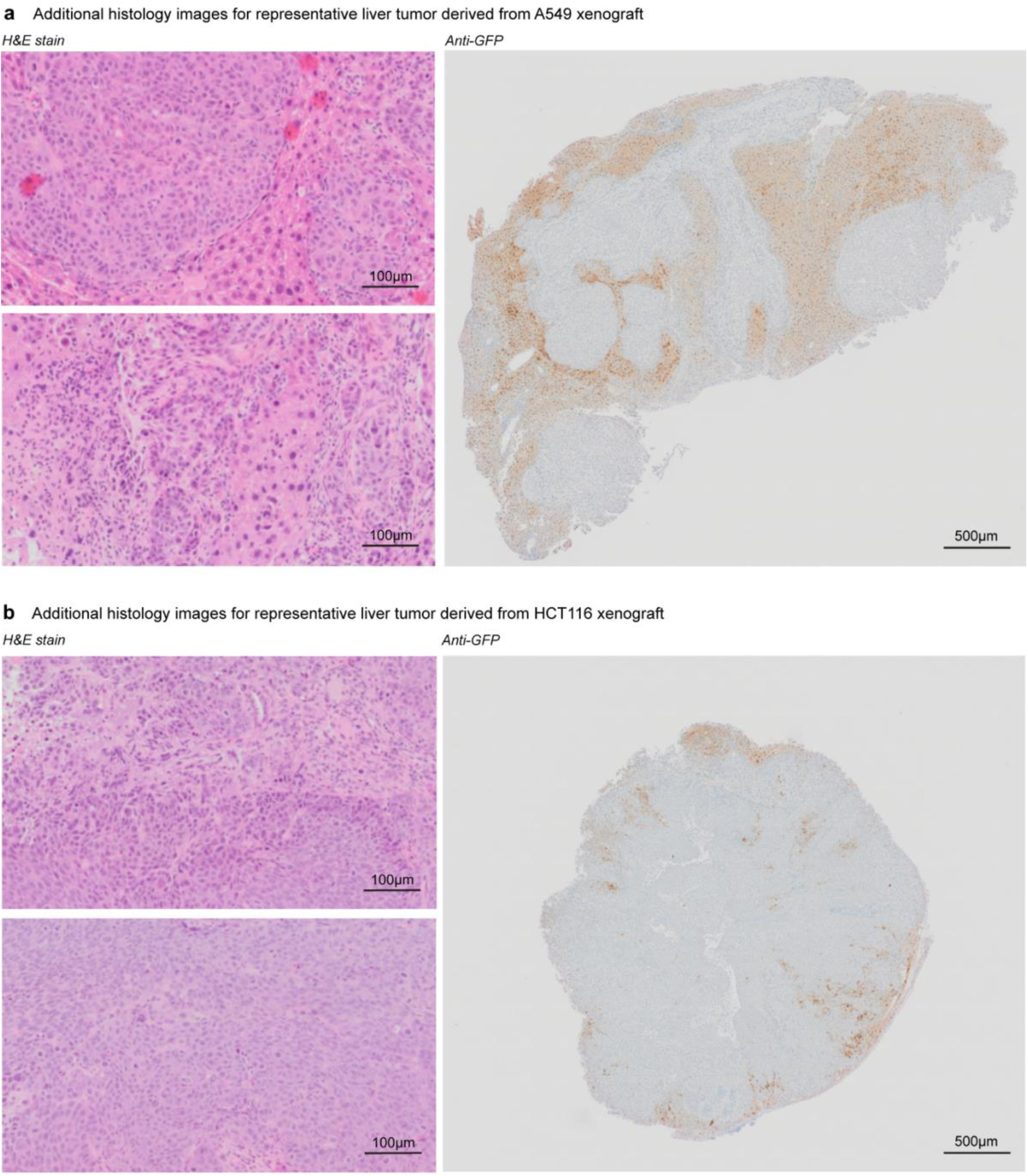
Additional histology images for liver xenografts modelling secondary liver cancer. H&E stain from representative regions of the tumor demonstrates tissue architecture. Anti-GFP stain shows location of eGFP delivery within the tumor. **(a)** liver tumor derived from A549 lung adenocarcinoma cells. **(b)** liver tumor derived from HCT116 colorectal carcinoma cells.

**Supplementary Figure 19:**
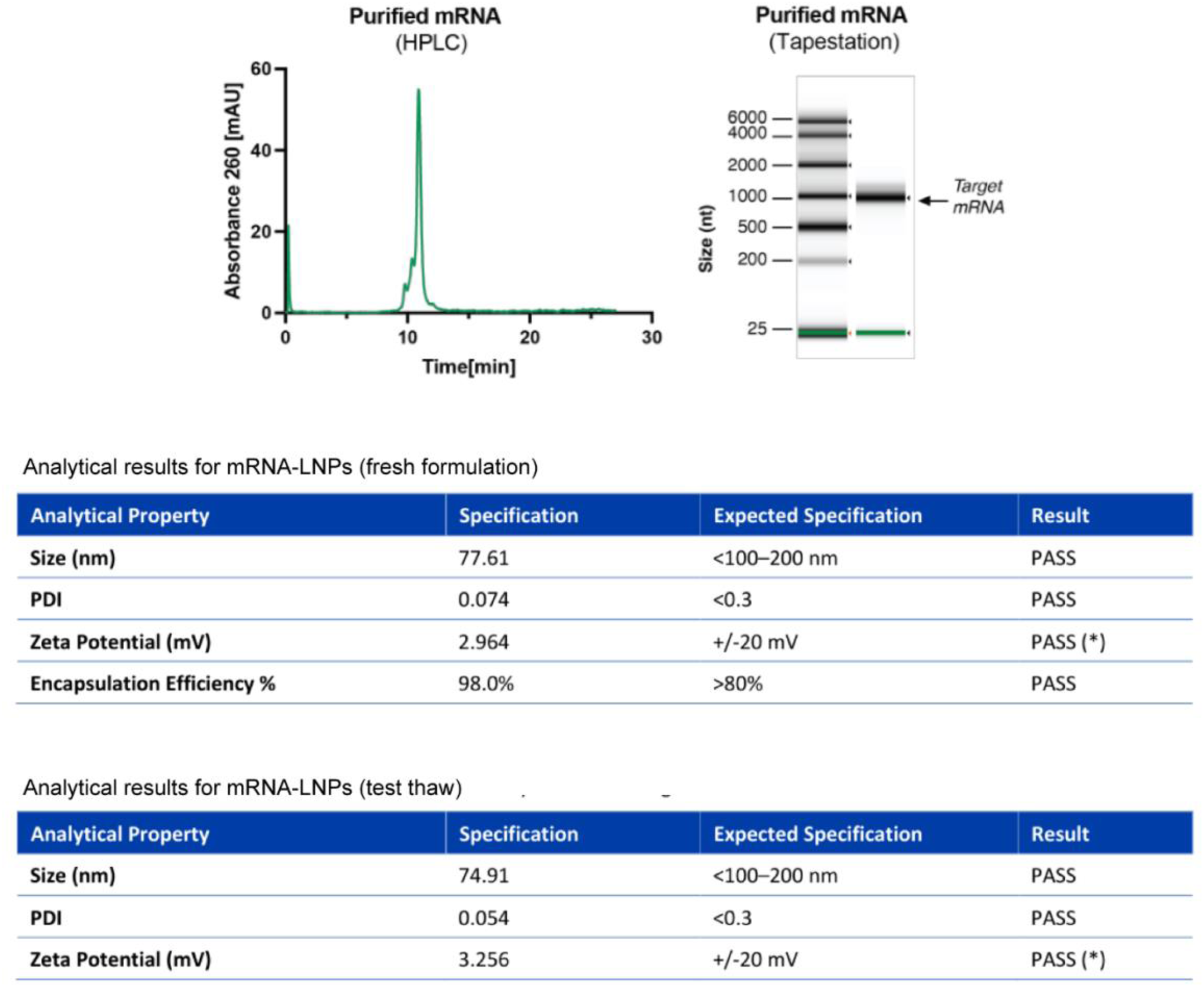
Excerpt from quality analysis of mRNA and mRNA-LNPs. Excerpt from a production report for a representative batch of eGFP mRNA and mRNA-LNPs, showing the expected size and purity of the synthetic mRNA, and the LNPs within specifications.

**Supplementary Table 1:** Genes differentially regulated in liver tissue 24 hours after mRNA-LNP injection. Supplementary Table 1 is supplied separately in .xlsx format.

**Supplementary Table 2:** Antibody details.

| Application | Antibody | Host | Supplier | Cat# | Dilution |
| --- | --- | --- | --- | --- | --- |
| Immunohistochemistry | Anti-GFP | Rabbit | Novus | NB600-308 | 1:1000 |
| Western blotting | Anti-GFP | Rabbit | Novus | NB600-308 | 1:2000 |
| Western blotting | Anti-rabbit IgG, highly cross-adsorbed, Alexa Fluor™ Plus 800 conjugate | Donkey | Thermo Fisher | A32808 | 1:10000 |

